# Synthetic Hybrid Receptors for Safer and Programmable T Cell Therapy

**DOI:** 10.64898/2026.01.22.701150

**Authors:** Maxwell G. Foisey, Julie Garcia, Xun Li, Xiaoyu Yang, Claire Hilburger, Chloe Thienpont, Alejandro Chaves-Martinez, Serkan Belkaya, Tina Truong, Iowis Zhu, Raymond Liu, Axel Hyrenius-Wittsten, Ricardo Almeida, Brian R. Shy, Greg M. Allen, Sarah Wyman, Kole T. Roybal

## Abstract

Engineered T cell therapies have achieved significant clinical success in hematological malignancies but remain largely ineffective in solid tumors. Overcoming this limitation requires strategies that enhance T cell function while avoiding systemic immune toxicities and pathological T cell states. Existing approaches typically rely on constitutive gene overexpression or suppression to augment potency or remodel the tumor microenvironment, but these strategies frequently lead to dysregulated immune activation and dose-limiting toxicity. Here, we present Hybrid Receptors (Hybrid-Rs), a modular receptor platform that integrates features of chimeric antigen receptors (CARs) and SyNthetic Intramembrane Proteolysis Receptors (SNIPRs) to couple antigen-dependent T cell activation with programmable gene regulation. Hybrid-Rs enable precise, context-dependent control of T cell potency, differentiation states, and conditional expression of secreted immunotherapeutic payloads with otherwise prohibitive toxicity. Hybrid-Rs are readily humanized and compatible with precision genome editing in primary human T cells, providing a direct and practical path to clinical translation.

## INTRODUCTION

Chimeric antigen receptor (CAR) T cell therapies have achieved remarkable clinical success in hematological malignancies, with CD19-directed CAR T cells producing objective response rates of up to 82% in patients with B cell lymphoma^1^. CARs redirect T cells to tumor-associated antigens and induce direct cytotoxicity through rapid, MHC-independent signaling analogous to the endogenous T cell receptor^2,3^. FDA-approved CARs also incorporate signaling chains from costimulatory receptors such as CD28 and 4-1BB for more robust T cell activation that improves the magnitude and persistence of anti-tumor activity^1,4^. Despite encouraging results in hematological malignancies, there are no FDA approved CAR T cell therapies for solid tumors, accounting for the majority of cancers (>90%). Thus, engineering next generation T cells that can effectively target and eliminate most cancers is critical to making cell therapies a pillar of cancer treatment^5^.

Efforts to improve CAR T cell performance in solid tumors have focused on receptor optimization and the addition of engineered functions, although relatively few new designs have advanced to clinical testing. Common strategies include constitutive co-expression of proinflammatory cytokines (“armored” CARs), transcription factors, or targeted gene knockouts to enhance effector function and persistence^6–13^. While there has been recent success in clinical trials with CAR T cells that constitutively produce IL-15 or IL-18^8,9^, not all immune stimulatory factors can be safely produced in an unregulated fashion as they can lead to systemic toxicity or T cell dysfunction, limiting therapeutic utility^14–17^.

Synthetic transcriptional receptor systems, such as synNotch and SyNthetic Intramembrane Proteolysis Receptors (SNIPRs), provide an alternative regulatory paradigm for therapeutic T cells^18,19^. Inspired by natural Notch signaling, these receptors undergo ligand-induced proteolytic cleavage to release an intracellular transcriptional regulator that drives antigen-dependent gene expression^18–20^. SynNotch and SNIPRs can be engineered to sense tumor-associated antigens similar to CARs, including cell surface antigens and soluble factors in the tumor environment, and upon ligation, initiate expression and production of therapeutic genes or payloads^19,21^. A broad range of cell intrinsic and extrinsic therapeutic payloads have been engineered into these antigen-inducible transcriptional circuits including transcription factors, cytokines, antibodies, toxins, adjuvants, and CARs^18–24^. A key feature of synNotch or SNIPR T cells is their ability to locally and precisely deliver potent immune stimulatory factors to tumors, improving efficacy and mitigating systemic toxicity potential.

Here, we introduce Hybrid Receptors (Hybrid-Rs), a new class of synthetic receptors that integrates the fast, cytotoxic signaling of CARs with the programmable gene-regulatory capacity of synNotch and SNIPRs in a single, compact architecture. Hybrid-Rs simultaneously activate T cells and induce customizable transcriptional programs upon antigen recognition, coupling immediate tumor cell killing with localized delivery of therapeutic payloads. Mechanistically, Hybrid-Rs signal at the plasma membrane in a CAR-like manner while also undergoing proteolytic cleavage to release a transcriptional regulator that initiates gene expression. We demonstrate that Hybrid-Rs can be engineered to target diverse tumor-associated antigens, incorporate multiple CAR signaling domains, and be fully humanized. Hybrid-R T cells exhibit robust antitumor activity across tumor models and enable controlled expression of potent efficacy-enhancing genes, including IL-12 and CARD11–PIK3R3, with improved safety in vivo. The compact, clinically compatible design of Hybrid-Rs facilitates integration via precision genome editing, providing a direct path toward clinical translation in both hematological and solid tumors.

## RESULTS

### Hybrid-Rs balance signaling and custom gene regulation in T cells

Receptor tyrosine kinases (RTKs) are responsible for a broad range of cellular functions^25^. A subset of RTKs undergo γ-secretase–mediated proteolysis^26^, however, unlike canonical Notch receptors, their intracellular domains (ICDs) are not restricted to nuclear translocation. Instead, RTK ICDs, such as that of Epharin type-B receptor 2, initiate kinase signaling and undergo proteolytic cleavage like Notch ^27,28^. This observation motivated us to design synthetic receptors capable of stimulating multiple responses, including T cell signaling and transcriptional regulation. If achieved, such a receptor could effectively elicit a potent native T cell response alongside synthetically programmed therapeutic functions (Fig. 1a, b left).

**Fig. 1:**
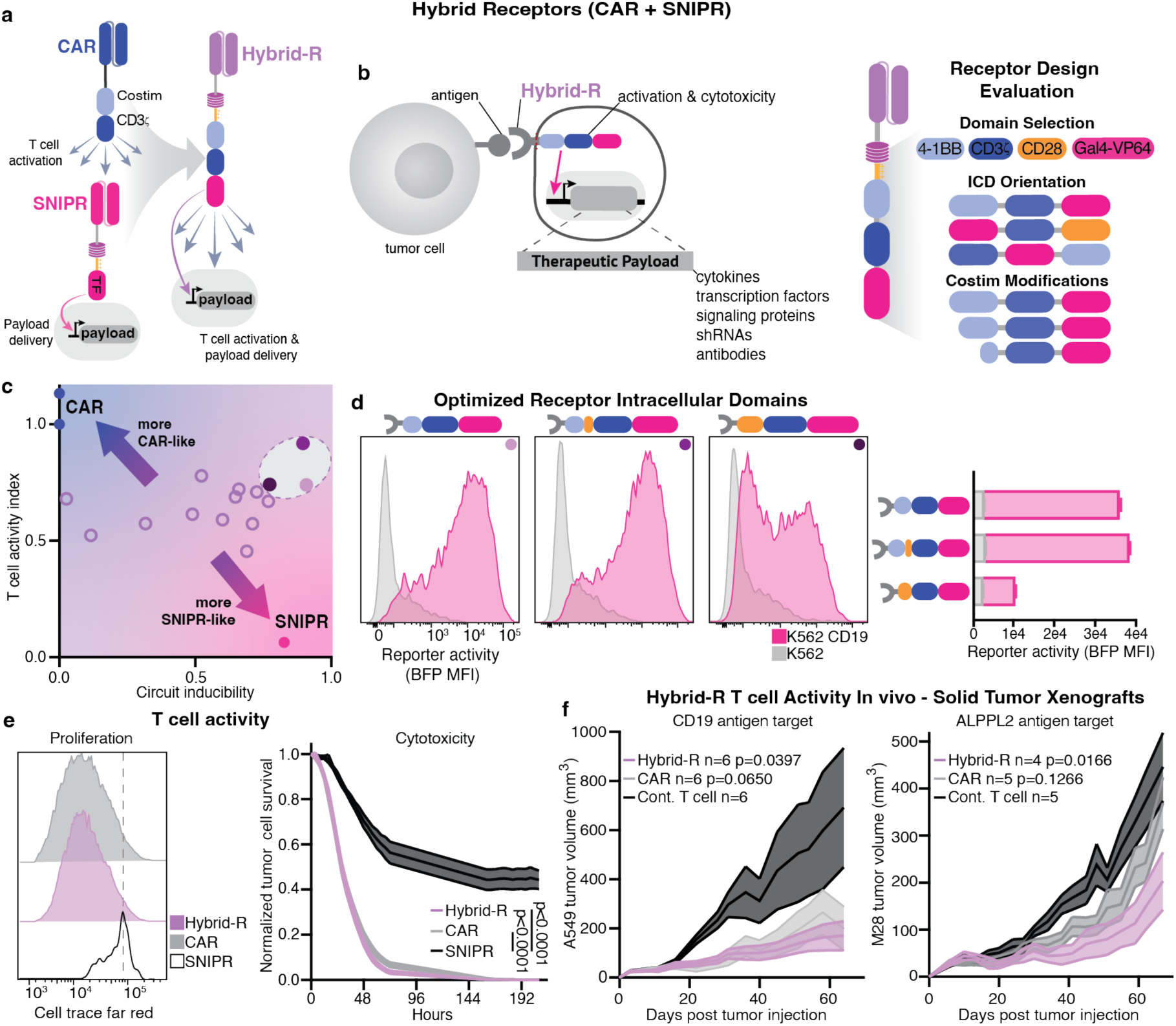
Hybrid Receptors (Hybrid-Rs) – synthetic receptors for programmed T cell activation and custom therapeutic gene regulation. **a**, Schematic showing Hybrid-R structure. **b**, Left, Schematic showing Hybrid receptor activity in primary human T cells. Right, Schematic showing Hybrid receptor optimization process. **c**, Normalized receptor activity in primary human T cells (representative of 3 experiments). T cell activity index is an aggregate score including surface marker expression, IL-2 production and cytotoxic capacity. Circuit inducibility is the normalized transcriptional circuit signal to noise ratio of stimulated and unstimulated Hybrid-R T cells. d, Left, Histograms depict optimized Hybrid-R mediated induction of BFP reporter transgene after 48 hours stimulation with CD19^+^ K562 target cells (representative of 3 experiments). Right, Quantification (MFI) of BFP reporter expression in Hybrid-R T cells (data from 3 independent donors). **e**, Left, cell trace far red dye dilution in SNIPR, Hybrid-R or CAR T cells after stimulation with CD19^+^ K562 target cells (representative of 2 experiments). Right, relative target cell lysis by anti-CD19 CARs, SNIPRs or Hybrid-Rs targeting CD19^+^ A549mKate-NLS cells measured by Incucyte live-cell imaging. Effector to target ratio 1:1. Data are total area normalized to untransduced control T cells (data from 2 independent donors n=2 each). Statistics calculated with two tailed unparied T tests. **f**, Left, Tumor volume of CD19^+^ A549 tumor bearing animals. Right, ALPPL2^+^ M28 tumor-bearing animals treated with engineered T cells. Statistics analyzed using one-way ANOVA with Dunnett’s correction comparing the area under the curve versus the untransduced control group over the course of the experiment. Mean SEM depicted.

To construct a synthetic receptor that couples T cell activation with inducible gene regulation, we defined a set of key domains drawn from second-generation CARs and SNIPRs that, in combination, could support both functions within a single receptor, termed a Hybrid-R. These domains include an antibody-based ligand-binding module, CAR-derived costimulatory and CD3ζ signaling elements, and SNIPR-derived proteolytic and transcriptional control components, including a cleavable hinge, transmembrane and juxtamembrane domains (JMD), and a transcriptional regulator ^19^. Guided by this framework, we then systematically rearranged intracellular domain (ICD) elements to identify Hybrid-R architectures that optimally balance proximal T cell signaling and inducible transcriptional activity in the context of an anti-CD19 scFv (Fig. 1b right). Primary human T cells expressing Hybrid-R variants were co-cultured with CD19⁺ K562 target cells for 48 hours, after which T cell activity and circuit inducibility were assessed (Fig. 1c, Extended data fig. 1). T cell activity was defined by IL-2 production, surface activation marker expression, and cytotoxicity, whereas circuit inducibility was quantified by basal and antigen-induced expression of a BFP transcriptional reporter (Fig. 1c, Extended data fig. 1a-f).

Receptors containing CD28-CD3ζ-Gal4-VP64 or 4-1BB-CD3ζ-Gal4-VP64 ICDs exhibited robust, antigen-dependent T cell activation but also displayed antigen-independent transcriptional activity (Extended data fig. 1a-f). In contrast, Hybrid-Rs with Gal4-VP64 positioned in the membrane-proximal ICD region, such as Gal4-VP64-4-1BB-CD3ζ, showed high-fidelity transcriptional induction but markedly reduced T cell activation (Extended data fig.1a–f), indicating a requirement for T cell–activating motifs to occupy membrane-proximal positions. Notably, a CD3ζ-Gal4-VP64 Hybrid-R exhibited partially balanced signaling and transcriptional outputs, suggesting that in some configurations costimulatory domains may be dispensable (Extended data fig.1a–f). Consistent with prior observations^29^, these data reinforced the requirement for membrane-proximal placement of T cell signaling domains to achieve effective downstream activation. We therefore shifted our focus toward improving Hybrid-R circuit fidelity through direct engineering of costimulatory domain sequences (Fig. 1b, right).

The linear signaling motifs of the CD28 and 4-1BB costimulatory domains are well characterized, and prior work has established a critical role for the JMD in transcriptional receptor function¹⁹. In the initial Hybrid-R designs, the ICD contained both the Notch-derived JMD required for transcriptional activation and the native JMDs of CD28 or 4-1BB, resulting in duplicated JMD elements that we reasoned were unnecessary and potentially detrimental to receptor performance. Because the essential signaling motifs of CD28 and 4-1BB are distal to the JMD, we tested whether N-terminal truncation of the costimulatory ICDs could reduce basal transcription while preserving T cell activation. Removing the costimulatory JMD sequences while retaining the Notch JMD was insufficient to suppress antigen-independent circuit activity (Extended data fig.1g). In contrast, progressive truncation of the costimulatory domain to retain only the core signaling motifs improved circuit fidelity and preserved potent T cell activity. (Fig. 1c,d; Extended data fig. 1g–i). These optimized Hybrid-Rs exhibited a more optimal balance between T cell activation and circuit inducibility than the original designs (Fig. 1c). Notably, a Hybrid-R incorporating a truncated 4-1BB domain together with the CD28 PYAP motif exhibited robust activity, demonstrating that custom-engineered signaling domains can be designed to precisely sculpt cellular responses. (Fig. 1c,d; Extended data fig. 1).

We selected a Hybrid-R incorporating a fully truncated 4-1BB–CD3ζ–Gal4-VP64 ICD for further study (Fig. 1c). Relative to SNIPRs, this optimized Hybrid-R exhibited stronger transcriptional circuit induction (Extended data fig. 1j). Consistent with prior reports that Notch receptor processing is enhanced by T cell activation^30^ the optimized Hybrid-R improved transcriptional output without compromising circuit fidelity. Direct comparison with CAR T cells revealed comparable proliferative capacity and cytotoxic activity, indicating that the optimized Hybrid-R does not impair core T cell functions (Fig. 1e).

To further contextualize these effects, we evaluated induction of native transcriptional pathways and T cell differentiation downstream of CAR, SNIPR, or the optimized Hybrid-R. Using an NFAT, NFκB, and AP-1 reporter Jurkat cell line^10^, we observed significantly greater activation of these pathways downstream of CAR and Hybrid-R relative to SNIPR (Extended data fig. 2a). Consistent with these findings, primary human Hybrid-R T cells stimulated with CD19⁺ K562 target cells displayed surface marker profiles indicative of an effector T cell phenotype, similar to CAR T cells (Extended data fig. 2b). Collectively, these data demonstrate that Hybrid-R integrates robust native T cell signaling with programmable transcriptional circuitry, enabling coordinated regulation of cellular activity and gene expression within a single receptor architecture.

To demonstrate the therapeutic potential of Hybrid-Rs, we next validated the optimized receptor architecture using multiple tumor-targeting scFvs. Upon stimulation with CD19-, ALPPL2-, or HER2-expressing K562 target cells, Hybrid-R T cells with corresponding scFvs retained both robust T cell activity and inducible circuit signaling (Extended data fig. 3a–d). To benchmark therapeutic performance against clinical comparators, including a CD19 CAR (Kymriah)^31^ and an ALPPL2 CAR^24^ we challenged NSG immunodeficient mice bearing solid tumors and treated them with either CAR or Hybrid-R T cells (Fig. 1f). Across solid tumor models targeting distinct tumor-associated antigens, Hybrid-R T cells conferred CAR-like control of tumor growth (Fig. 1f). These data indicate that the optimized Hybrid-R architecture is compatible with diverse tumor targets and binding domains, and that its modular design enables systematic tuning for broad therapeutic applications.

### Hybrid-Rs control therapeutic gene programs in vivo

Next-generation CAR T cell therapies increasingly incorporate immunomodulatory payloads that can be classified as T cell–extrinsic or T cell–intrinsic^7–10^. Whereas extrinsic payloads act in cis or trans on surrounding cells, intrinsic payloads directly reprogram T cell state and function. To evaluate the therapeutic potential of Hybrid-Rs, we coupled a potent T cell–intrinsic immunomodulatory program to Hybrid-R circuitry.

We selected the CARD11–PIK3R3 (CP) fusion, a strong amplifier of NF-κB and AP-1 signaling identified in an in vivo pooled screen of human T cell lymphoma mutations (Fig. 2a)^10^. Although CP fusion enhances T cell potency downstream of TCR or CAR engagement, unregulated expression in the context of an intact endogenous TCR repertoire induces toxic alloreactivity in vivo¹⁰. We therefore evaluated Hybrid-R–mediated regulation of CP fusion expression in a stringent syngeneic tumor model using hCD19⁺ B16 melanoma tumors, without lymphodepletion or preconditioning. Consistent with prior reports, unregulated CP fusion expression enhanced CAR T cell potency and resulted in tumor control accompanied by marked toxicity, with nearly all treated animals succumbing to treatment-associated adverse effects (Fig. 2b,c; Extended data fig. 4a,b). In contrast, Hybrid-R–mediated control of CP fusion expression enabled uniform tumor control in all treated mice (6/6) without observable toxicity (0/6) (Fig. 2b,c). These data demonstrate that Hybrid-R circuits can safely deploy highly potent T cell–intrinsic payloads, thereby expanding the therapeutic window of engineered T cell therapies.

**Fig. 2:**
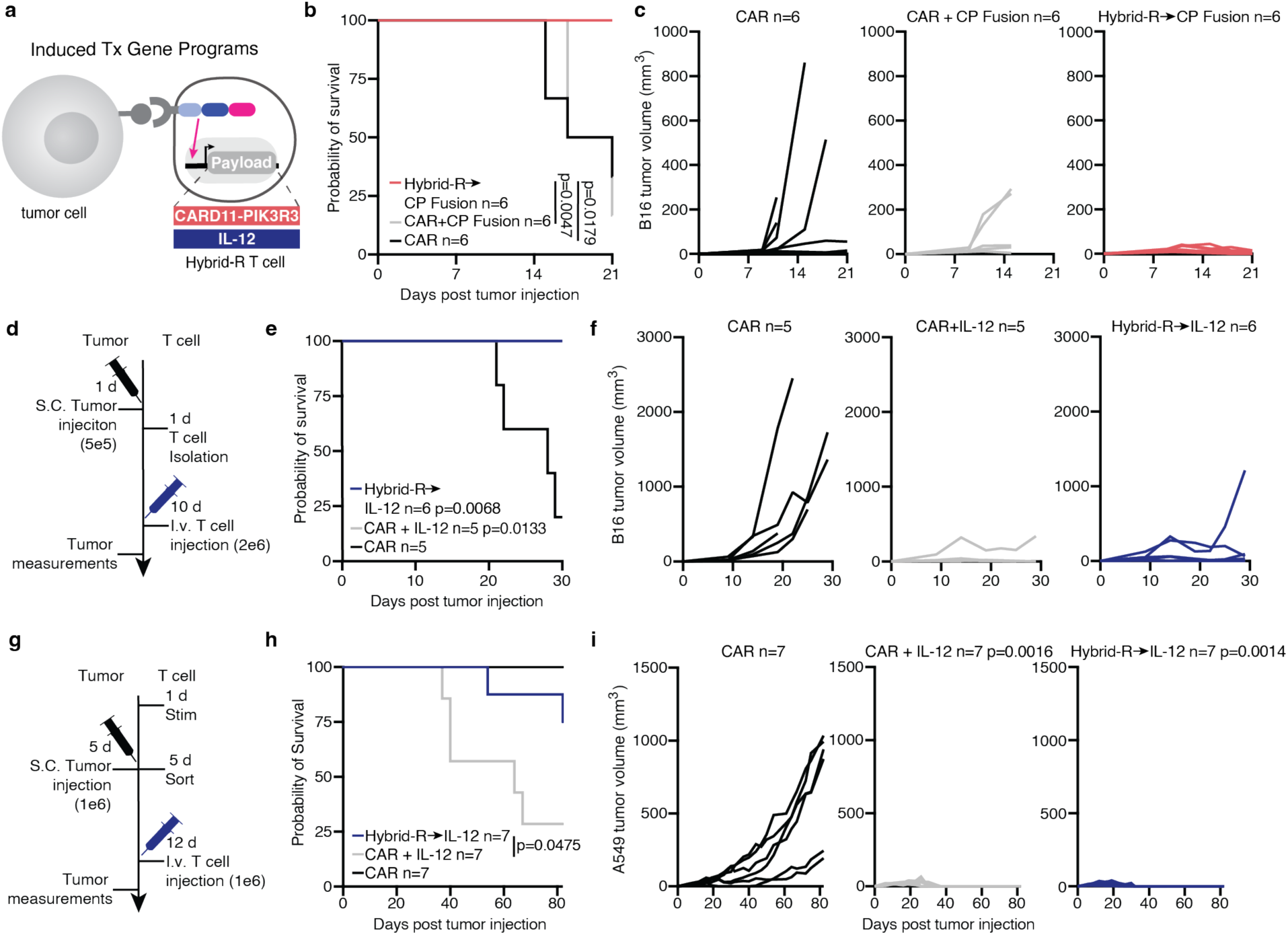
Hybrid-Rs safely enhance T cell potency and extend survival by regulating therapeutic gene programs. **a**, Schematic of Hybrid-R→Payload T cell activity. **b**, Survival of hCD19^+^ B16 tumor bearing animals after treatment with 4.0e6 engineered T cells 9 days after tumor innoculation. P values calculated using log-rank Mantel-Cox survival. **c**, Tumor volumes from hCD19^+^ B16 tumor-bearing syngeneic animals treated with Hybrid-R→CP Fusion T cells. Lines indicate individual mice. **d**, Schematic of in vivo study. **e**, Survival of hCD19^+^ B16 tumor bearing syngeneic animals treated with 2.0e6 Hybrid-R→IL-12 T cells 9 days after tumor innoculation. P values calculated using log-rank Mantel-Cox survival. **f**, Tumor volumes of hCD19^+^ B16 tumor-bearing syngeneic animals treated with Hybrid-R→IL-12 T cells. Lines indiciate individual mice. **g**, Schematic of in vivo study. **h**, Survival of CD19^+^ A549 tumor bearing NSG animals treated with 1.0e6 Hybrid-R→IL-12 T cells 7 days after tumor innoculation. P values calculated using log-rank Mantel-Cox survival. **i**, Tumor volumes of CD19^+^ A549 tumor-bearing NSG animals. Tumor volume statistics analyzed using one-way ANOVA with Dunnett’s correction comparing the area under the curve versus the CAR control group over the course of the experiment. Lines indicate individual mice.

Having established Hybrid-R–mediated control of potent T cell–intrinsic payloads, we next assessed its capacity to regulate T cell–extrinsic effectors. We selected the pleiotropic immunostimulatory cytokine IL-12 as a model cell extrinsic payload to place under Hybrid-R control (Fig. 2a). IL-12 has demonstrated strong anti-tumor activity through enhancement of T cell effector function, interferon-γ signaling, and remodeling of the tumor microenvironment^32,33^. however, its clinical utility has been limited by dose-dependent toxicities when not tightly regulated^11,33–36^. Using the same syngeneic hCD19⁺ B16 melanoma model without lymphodepletion, both Hybrid-R→IL-12 and CAR+IL-12 T cells significantly improved tumor control and survival (Fig. 2d–f). To further evaluate safety and translational relevance, we next tested Hybrid-R–mediated IL-12 delivery in human T cells in NSG mice bearing CD19⁺ A549 solid tumors (Fig. 2g). In this setting, Hybrid-R→IL-12 T cells achieved durable tumor eradication while limiting treatment-associated toxicity (Fig. 2h,i; Extended data fig. 4c). Similar results were observed in a CD19⁺ A375 solid tumor model, in which Hybrid-R→IL-12 T cells consistently improved tumor control while maintaining favorable safety profiles (Extended data fig. 4d–f).

### Benchmarking Hybrid-R circuits against state-of-the-art T cell therapies

Transcriptional control of therapeutic payloads downstream of CAR or TCR signaling has previously been deployed clinically to treat solid tumors. Armored CAR T cells typically leverage native or engineered transcriptional response elements (REs), including NFAT, NFκB, and AP-1, to induce transgene expression following TCR or CAR activation^33,36,37^. However, these approaches rely on endogenous T cell signaling pathways. In contrast, Hybrid-R employs orthogonal, synthetic transcriptional control decoupled from native activation programs. We therefore benchmarked Hybrid-R against RE-based armored CAR platforms.

Using a BFP reporter to quantify circuit behavior following tumor antigen stimulation, Hybrid-R exhibited high-fidelity, antigen-dependent transgene induction relative to RE-based circuits (Fig. 3a; Extended data fig. 5a). Hybrid-R also showed increased signal amplitude and fold induction compared with NFAT- and NFκB-gated systems (Fig. 3b; Extended data fig.5b). Notably, upon removal of antigen-positive target cells, Hybrid-R circuits returned to baseline activity, whereas RE-based circuits remained active (Fig. 3a; Extended data fig. 5a). Moreover, Hybrid-R circuits remained inactive under TCR stimulation alone, confirming that transgene expression is restricted to Hybrid-R ligation and synthetic signaling rather than general T cell activation (Fig. 3a; Extended data fig. 5a).

**Fig. 3:**
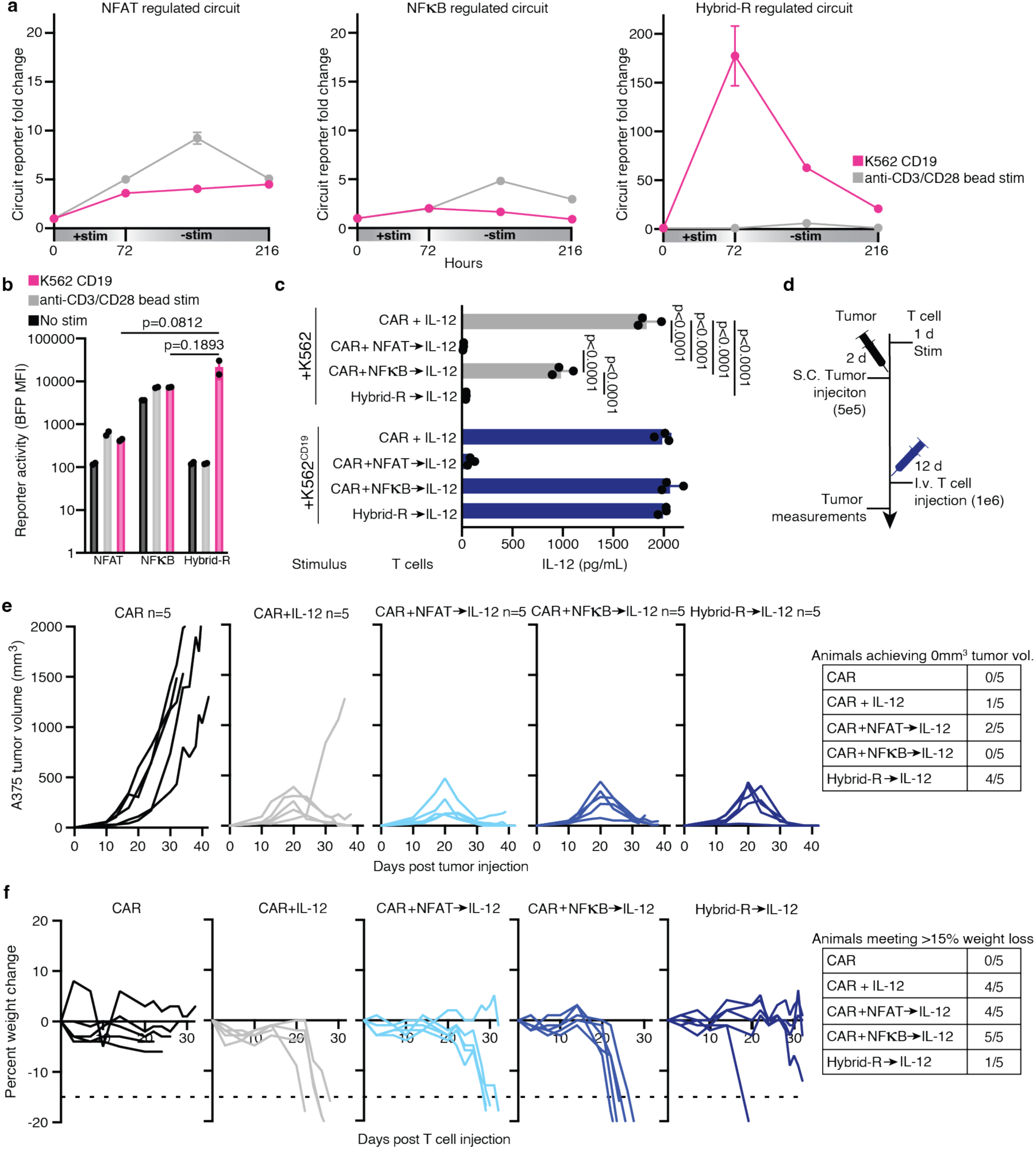
Hybrid-R circuits match functional performance and improve safety relative to T cell signaling response element–based regulation. **a**, Left, Fold change in circuit reporter activity (MFI) from anti-CD19 4x NFAT or 5x NFkB armored CAR or Hybrid-R T cells in presence of stimuli relative to resting NFAT, NFkB or Hybrid-R T cells. Data from 2 independent donors. **b**, BFP reporter MFI values from engineered T cells at 72 hours. (data from 2 independent donors). **c**, Quantification of human IL-12 in cell culture supernatant is given after coculture for 72 hours with CD19^+^ K562 target cells (n=3 technical replicates). Statistics calculated with one-way ANOVA with Šídák’s multiple comparisons test. Mean SEM is depicted. **d**, Schematic of in vivo study. **e**, Tumor volume from CD19^+^ A375 solid tumor-bearing NSG animals treated with 1.0e6 IL-12 producing T cells. **f**, CD19^+^ A375 tumor bearing animal percent weight change. Lines indicate individual mice, 15% weight loss represents euthanasia criteria.

Because IL-12 has previously been placed under RE control in clinical armored CAR T cells^36^, we next benchmarked Hybrid-R–mediated IL-12 regulation against RE-based approaches. Hybrid-R→IL-12 T cells exhibited superior control over cytokine production compared with constitutive or RE-mediated expression, producing high IL-12 levels only upon engagement with antigen-positive target cells (Fig. 3c; Extended data fig.5c). Given that interferon-γ contributes to IL-12–associated toxicity³⁴, we assessed downstream cytokine induction in vitro and observed that Hybrid-R–regulation of IL-12 limited non-specific interferon-γ secretion (Extended data fig. 5d).

To directly compare in vivo therapeutic performance and toxicity, we evaluated Hybrid-R→IL-12 T cells alongside NFAT- and NFκB-gated armored CAR T cells in the CD19⁺ A375 solid tumor model (Fig. 3d). At a failure dose, CAR-only T cells did not control tumor growth (Fig. 3e), while IL-12 armored CAR T cells showed partial tumor control, but were associated with substantial toxicity and weight loss across RE configurations (Fig. 3e,f). In contrast, 4 of 5 mice treated with Hybrid-R→IL-12 T cells achieved tumor control below the limit of detection, with only 1 of 5 animals exhibiting severe weight loss (Fig. 3f). Together, these data demonstrate that Hybrid-R provides improved control over genetically encoded immunotherapeutic payloads and favorably benchmarks against state-of-the-art transcriptional regulators.

### Hybrid-R are amenable to clinical translation

We have thus far demonstrated the activity and safety of Hybrid-Rs incorporating the Gal4–VP64 transcriptional activator; however, the use of non-human components introduces potential immunogenicity that may limit clinical translation. To generate a more clinically tractable Hybrid-R system, we replaced Gal4–VP64 with a human-derived HNF1α–p65 transcriptional activator (Fig. 4a)^19,38^. Upon antigen stimulation, this humanized Hybrid-R exhibited circuit strength comparable to the original configuration (Fig. 4b,c) while retaining robust T cell effector function, as evidenced by effective control of target cell growth in vitro (Fig. 4d).

**Fig. 4:**
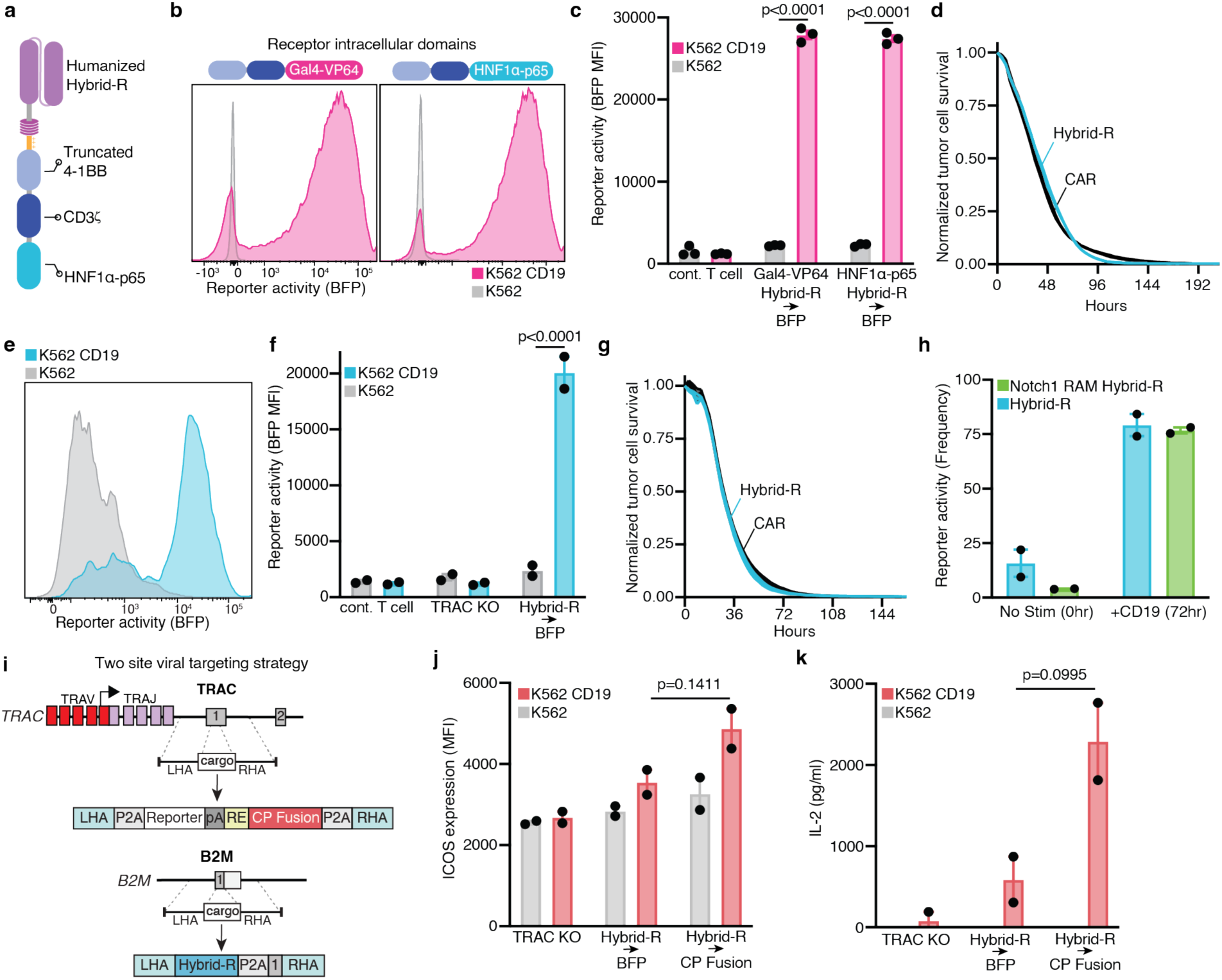
Humanized Hybrid-R circuits are amenable to site-specific knock-in and exhibit robust activity. **a**, Schematic of humanized Hybrid-R. **b**, Histograms depict antigen dependent induction of BFP reporter in lentiviral Hybrid-R T cells in response to 72 hours stimulation with CD19^+^ K562 target cells (representatvie of 2 experiments). **c**, Quantification (MFI) of BFP reporter expression (n=3 technical replicates). Statistics calculated with two-way Anova with Šídák’s multiple comparisons test. **d**, Relative target cell lysis by anti-CD19 Hybrid-Rs targeting CD19^+^ A549mKate-NLS cells measured by Incucyte live-cell imaging. E:T=1:2. Data are total area normalized to untransduced control T cells (n=2 technical replicates). **e**, BFP reporter gene expression in Hybrid-R two-site targeted knock-in primary human T cells 96 hours after stimulation with CD19^+^ K562 target cells (representative of 3 experiments). **f**, Quantification (MFI) of BFP reporter expression. Statistics calculated with one-way ANOVA with Šídák’s multiple comparisons test. Mean SEM is depicted. **g**, Relative target cell lysis by anti-CD19 Hybrid-Rs targeting CD19^+^ A549mKate-NLS cells measured by Incucyte live-cell imaging. E:T=1:4. Data are total area normalized to TRAC knockout control T cells. (data from 2 independent donors n=3 each). **h**, Quantification of BFP reporter transgene expression (frequency of mCherry^+^ T cells) when under optimized Hybrid-R or Notch1 JMD+RAM Hybrid-R (data from 2 independent donors n=3 each). **i**, Schematic of two site Hybrid-R→CP Fusion circuit targeting strategy. **j**, Surface marker ICOS expression (MFI) on engineered T cells 72 hours after stimulation with CD19^+^ K562 target cells **k**, Quantification of IL-2 in cell culture supernatant after coculture for 96 hours with CD19^+^ K562 target cells via ELISA (data from 2 independent donors n=3 each). Statistics calculated with one-way ANOVA with Šídák’s multiple comparisons test. Mean SEM is depicted.

To further advance Hybrid-Rs toward clinical translation, we adapted the receptor and circuit architecture for targeted genomic integration using CRISPR–Cas9. We implemented a two-site targeting strategy at the TRAC and B2M loci^39^, enabling stable and coordinated expression of the Hybrid-R and transcriptional circuit (Extended data fig.6a). Using dual AAV-delivered homology-directed repair templates, we achieved efficient dual knock-in in primary human T cells (Extended data fig. 6b). Upon co-culture with CD19⁺ target cells, gene-edited Hybrid-R T cells exhibited robust antigen-dependent circuit induction (Fig. 4e,f) and potent target cell–directed cytotoxicity (Fig. 4g).

In some configurations, targeting the transcriptional circuit to the TRAC locus increased basal circuit activity. Given prior reports that Notch receptor processing can be modulated by JMD selection (Notch1 versus Notch2) and by inclusion of the Notch-derived RBPJκ-associated molecule (RAM) domain^19,40^, we incorporated these elements into the Hybrid-R design. These modifications reduced basal circuit activity (Fig. 4h; Extended data fig.6c) but also attenuated maximal transcriptional output (Extended data fig. 6d,e), while T cell activation profiles relative to the original Notch2 JMD-based Hybrid-R were preserved (Extended data fig. 6f). We additionally tested whether basal activity arose from HNF1α response element (RE) activity at the TRAC locus by varying the number of HNF1α binding sites. While basal induction was unaffected, transcriptional output could be titrated without altering receptor architecture (Extended data fig. 6g), demonstrating tunable circuit behavior following genomic integration.

We next evaluated humanized two-site Hybrid-Rs regulating the CP Fusion. Efficient dual knock-in at the TRAC and B2M loci was observed (Fig. 4i, Extended data fig. 7a,b), and following antigen stimulation, CP Fusion expression was confirmed by reporter induction alongside increased IL-2 production and ICOS expression, consistent with expected functional outputs (Fig. 4j,k; Extended data fig. 7c–e)^10^.

To simplify manufacturing and mitigate concerns associated with multi-site targeting, we consolidated the Hybrid-R and transcriptional circuit into a single homology-directed repair template targeted to the TRAC locus (Fig. 5a). Using this single-vector configuration, we evaluated Hybrid-R activity against the clinically relevant solid tumor antigen B7-H3^41–44^. Efficient knock-in was achieved in primary human T cells (Extended data fig.8a), and upon co-culture with tumor cells expressing varying levels of B7-H3, Hybrid-R T cells exhibited high-fidelity circuit induction and potent cytotoxicity (Fig. 5b–d; Extended data fig.8b; Extended data fig. 9).

**Fig. 5:**
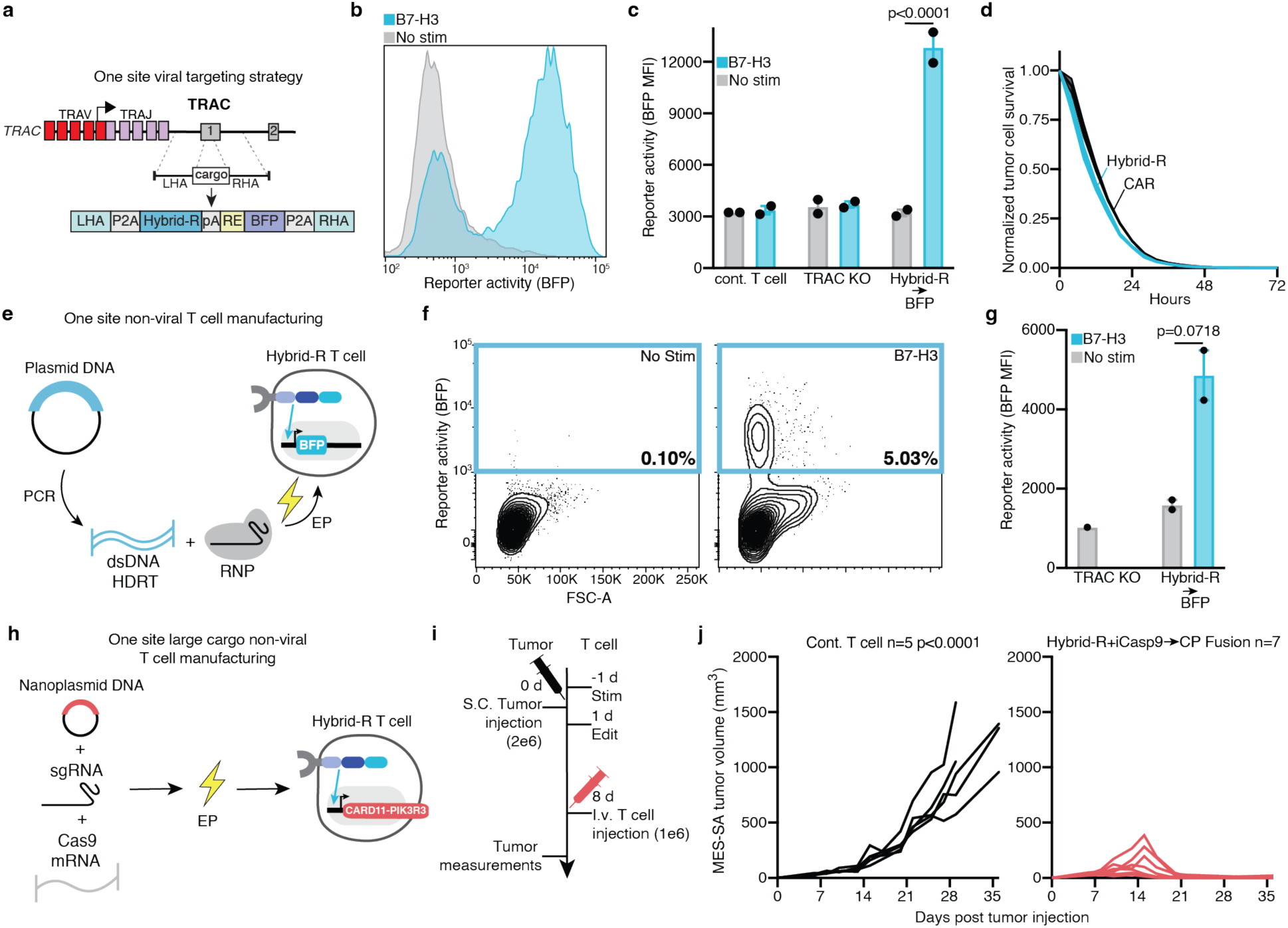
Single-site TRAC knock-in Hybrid-R circuits preserve function and support simplified manufacturing. **a**, Schematic of one-site viral Hybrid-R targeting strategy. **b**, BFP reporter expression in Hybrid-R knock-in primary human T cells 72 hours after stimulation with B7-H3^+^ A375 target cells. **c**, Quantification (MFI) of BFP reporter expression after 72 hours with B7-H3^+^ A549 target cells. (data from 2 independent donors n=3). **d**, Relative target cell lysis by anti-B7-H3 Hybrid-Rs targeting B7-H3^+^ MES-SAmKate-NLS cells measured by Incucyte live-cell imaging. Effector to target ratio is 1:4. Data are total area normalized to TRAC knockout T cells (n=3 technical replicates). **e**, Schematic of one-site non-viral Hybrid-R targeting strategy. **f**, Contour plots depict antigen dependent induction of BFP reporter in Hybrid-R T cells in response to 72 hours stimulation with B7-H3^+^ MES-SA target cells. Representitive of 2 biological replicates. **g**, Quantification (MFI) of BFP reporter expression (n=2 biological replicates). Statistics calculated with one-way ANOVA with Šídák’s multiple comparisons test. Mean SEM is depicted. **h**, Schematic of one-site non-viral targeting strategy. **i**, Schematic of in vivo study. **j**, Tumor volumes from B7-H3^+^ MES-SA tumor-bearing animals treated with 1.0e6 engineered T cells 8 days after tumor innoculation. Lines indicate individual mice. Tumor volume statistics analyzed using one-way ANOVA with Dunnett’s correction comparing the area under the curve versus the control group over the course of the experiment.

Because AAV cargo limitations constrain single-locus delivery of both receptor and payload, we adapted the single-vector Hybrid-R design for non-viral knock-in (Fig. 5e). Using double-stranded DNA and Cas9 ribonucleoprotein targeting the TRAC locus, we achieved efficient integration (15.7%) in primary human T cells (Extended data fig.8c). Upon co-culture with B7-H3⁺ MES-SA target cells, a human uterine sarcoma cell line, the non-virally manufactured Hybrid-R T cells exhibited antigen-dependent circuit induction with minimal basal activity (Fig. 5f,g). To further push toward clinical manufacturing compatibility, we generated a large-cargo, non-viral Hybrid-R vector encoding the receptor, an inducible payload, and an inducible Caspase9 (iCasp9) safety switch^45^, totaling 8.3 kb including homology arms. Using nanoplasmid DNA and Cas9 mRNA, we achieved efficient TRAC-targeted knock-in at lab-scale (Fig. 5h; Extended data fig. 8d). In NSG mice bearing B7-H3⁺ MES-SA tumors, Hybrid-R T cells controlled or eliminated tumors, whereas control T cells did not (Fig. 5i,j).

To assess clinical manufacturing compatibility at scale, we electroporated 5×10⁷ primary human T cells using the CTS Xenon system (Fig. 6a) and achieved efficient TRAC-targeted knock-in (9.62%) (Fig. 6b, Extended data fig. 10a). These cells effectively cleared B7-H3⁺ targets in vitro (Fig. 6c), and activation of the iCasp9 safety switch selectively depleted vector-positive cells (Fig. 6d). In vivo, Hybrid-R T cells conferred durable tumor control and significant survival benefit across dosing conditions (Fig. 6e-g). Together, these data establish a humanized Hybrid-R circuit, inserted into TRAC, manufactured with clinically viable methods that shows robust anti-tumor efficacy and contains a kill-switch to mitigate potential toxicities.

**Fig. 6:**
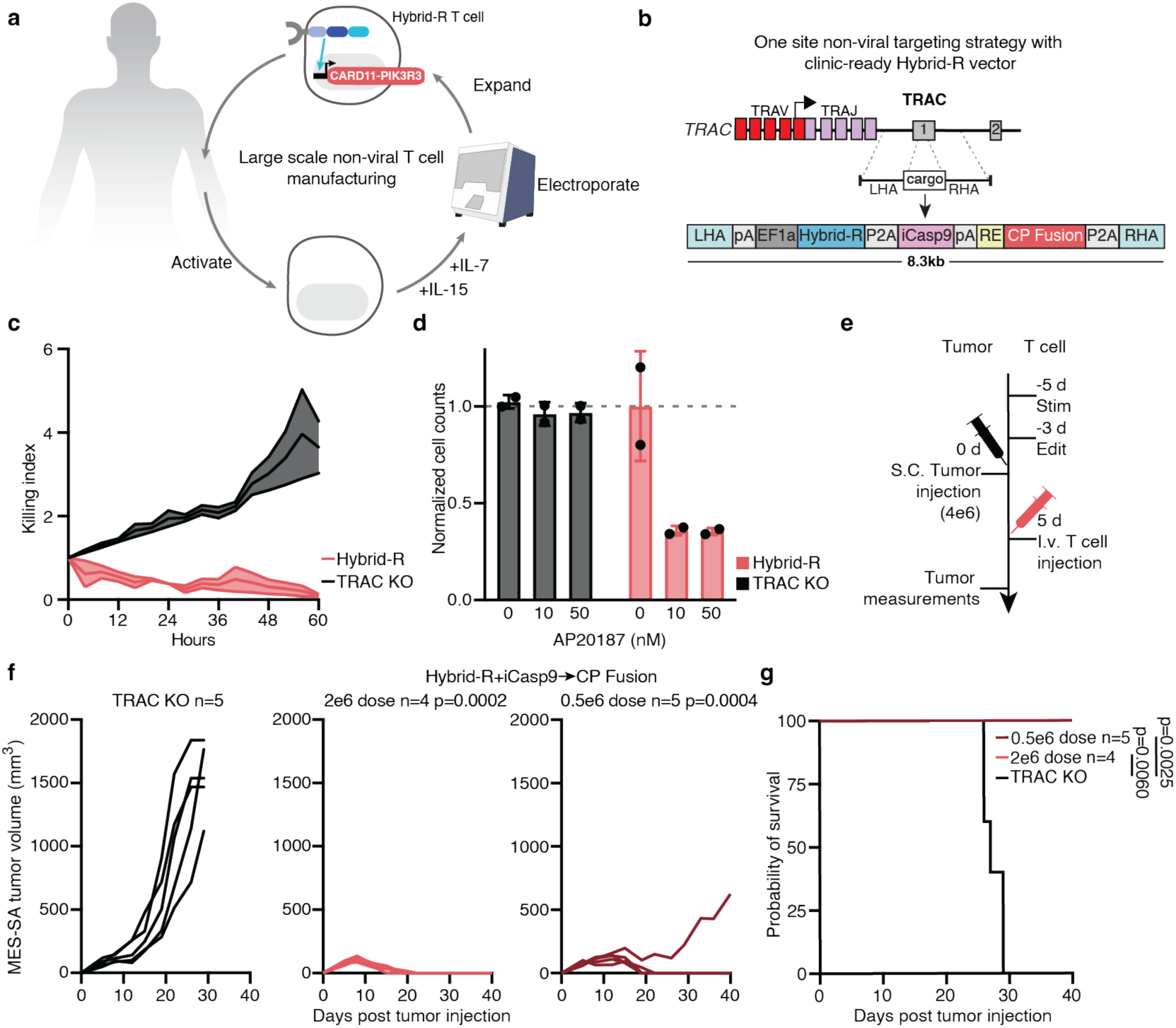
Clinical-candidate B7-H3 Hybrid-R→CARD11 Fusion T cells exhibit in vitro and in vivo safety and efficacy. **a**, Schematic of GMP compatible manufacturing strategy. **b**, Schematic of one-site non-viral targeting strategy. **c**, Relative target cell lysis by anti-B7-H3 Hybrid-Rs targeting B7-H3^+^ MES-SAmKate-NLS cells measured by Incucyte live-cell imaging. Effector to target ratio is 1:8. Data are total area normalized to first timepoint (n=3 technical replicates). **d**, Normalized T cell counts after 48 hours incubation with AP20187 (n=2 technical replicates). **e**, Schematic of in vivo study. **f**, Tumor volumes from B7-H3^+^ MES-SA tumor-bearing NSG animals treated with engineered T cells 5 days after tumor innoculation. Lines indicate individual mice.Tumor volume statistics analyzed using one-way ANOVA with Dunnett’s correction comparing the area under the curve versus the control group over the course of the experiment. **g**, Survival of B7-H3^+^ MES-SA tumor bearing NSG animals. P values calculated using log-rank Mantel-Cox (survival).

## DISCUSSION

Hybrid-R circuits enable a balance between efficacy and safety in engineered T cell therapies by integrating potent T cell activation with programmable, antigen-dependent regulation of therapeutic gene expression. Through synthetic transcriptional signaling, Hybrid-Rs decouple payload control from endogenous activation pathways, allowing T cells to mount robust anti-tumor responses while restricting therapeutic delivery or cellular enhancement to the tumor context. This coordinated control supports maximally effective immune activity alongside localized and tunable therapeutic intervention. The modularity of Hybrid-R design, combined with its amenability to humanization and site-specific genome integration in primary human T cells, provides a clinically viable framework for advancing next-generation cell therapies.

Beyond oncology, our systematic assembly and evaluation of synthetic receptor components highlight the broader potential of Hybrid-Rs to be precisely calibrated for additional disease settings. For example, Hybrid-Rs engineered to promote proliferation or persistence without cytotoxicity could enable T cells to function as self-sustaining, programmable drug delivery vehicles in inflammatory or degenerative conditions. More generally, the ability to tailor signaling and transcriptional outputs positions Hybrid-Rs as versatile tools for interrogating fundamental biological processes that depend on coordinated multicellular behavior. Deploying Hybrid-Rs across engineered immune cell subsets, such as macrophages and T cells, to transmit and receive user-defined signals could enable the coordinated control of therapeutic multicellular assemblies in vitro and in vivo.

Realizing these broader applications requires receptor–circuit systems that are not only functionally versatile but also compatible with clinical translation. In this regard, our work demonstrates a clear path from synthetic biology prototype to clinically translatable receptor–circuit systems. We advanced Hybrid-Rs from initial designs to single-site, genome-integrated vectors compatible with diverse manufacturing strategies and achieved non-viral integration of large payloads up to 8.3 kb in primary human T cells. As genome editing efficiencies continue to improve, even larger and more complex constructs may become feasible, enabling the coordinated deployment of multiple synergistic immunomodulatory programs under Hybrid-R control^46,47^. Together, these advances establish Hybrid-Rs as a versatile addition to the mammalian synthetic biology toolkit, with the potential to broaden the scope and impact of engineered cell therapies.

## Acknowledgments

K.T.R. is supported by the Parker Institute for Cancer Immunotherapy, the CRI Lloyd J. Old STAR Award and the NIH Director’s New Innovator Award (DP2 CA239143). M.G.F. is the recipient of a Parker Institute for Cancer Immunotherapy Scholar Fellowship.

## Contributions

K.T.R and M.G.F. conceived and designed the initial project, performed experiments, interpreted data and wrote the manuscript. J.G., A.W., I.Z, R.L., designed, performed and interpreted initial experiments. X.L., X.Y. designed, performed and interpreted experiments relating to two-site and one-site knock-in Hybrid-R T cells. T.T., X.Y., and B.S., designed performed and interpreted experiments relating to the GMP-compatible manufacturing of Hybrid-R T cells. C.H., C.T., A.C.M., and S.B., designed, performed and interpreted experiments relating to the evaluation of these cells in vitro and in vivo. G.M.A., R.A., B.S. supervised studies. S.W. and X.L. interpreted data and reviewed the manuscript.

## Data availability

All data associated with this study are present in the manuscript or its Supplementary Information Files

## Competing interests

M.G.F., X.L., J.G., A.W., R.L, I.Z., and K.T.R. have patents for Hybrid-R receptors or variants or immunotherapeutic payloads. J.G. is an employee of Moonlight Biotherapeutics. R.L. is an employee of Dispatch Biotherapeutics. K.T.R. is a co-founder of and stockholder in Arsenal Bio, Dispatch Biotherapeutics and Moonlight Bio. M.G.F., K.T.R., and X.L. are co-inventors on a provisional patent application that incorporates discoveries described in this manuscript.

## Description of Supplementary Materials

**Supplementary Table 1:** Plasmid names and T cell donor information for each experiment described.

**Supplementary Table 2:** Description, part, and amino acid and DNA sequences for all plasmids described.

**Supplementary Table 3:** Description and sourcing of genetic components used.

**Supplementary Table 4:** List of antibodies used.

**Supplementary Table 5:** List of sgRNAs used.

## Methods References

1. Graf, R., Li, X., Chu, VT., Rajewsky, K. sgRNA Sequence Motifs Blocking Efficient CRISPR/Cas9-Mediated Gene Editing. *Cell Rep*. **26**, 1098-1103 (2019).

2. Li, X. et al. “Precise CRISPR-Cas9 gene repair in autologous memory T cells to treat familial hemophagocytic lymphohistiocytosis.” *Sci. Immunol.* **9** (2024).

---

**Extended data fig. 1:**
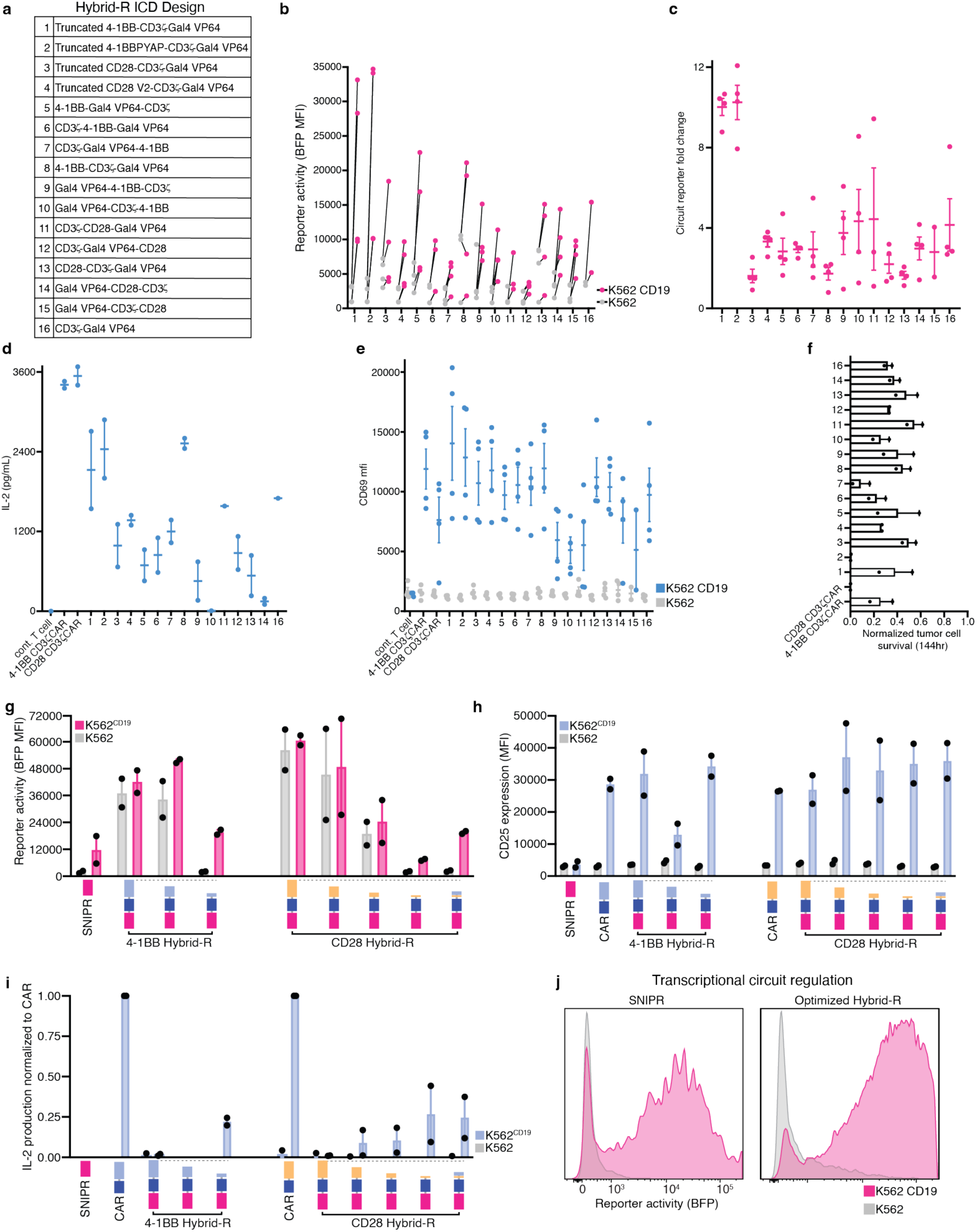
Hybrid-R design evaluation in primary human T cells. **a**, Schematic of Hybrid-R variants. **b**, BFP reporter gene expression (MFI) in Hybrid-R primary human T cells 48 hours after stimulation with CD19^+^ K562 target cells (data from 4 independent donors). **c**, Fold change in BFP reporter in primary human Hybrid-R T cells 48 hours after stimulation with CD19^+^ K562 target cells (data from 4 independent donors). **d**, Quantification of IL-2 production in cell culture supernatant after coculture for 48 hours with CD19^+^ K562 target cells (data from 2 independent donors). **e**, Quantification of CD69 surface expression (MFI) in Hybrid-R T cells 48 hours after stimulation with CD19^+^ K562 target cells (data from 4 independent donors). **f**, Quantification of total area of remaining by CD19^+^ A549mKate-NLS target cells at 144 hours after coculture with Hybrid-R T cells measured by Incucyte live-cell imaging. Effector to target ratio is 1:2. Data here are total area normalized to untransduced T cells (data from 2 independent donors). **g**, BFP reporter gene expression (MFI) 48 hours after stimulation with CD19^+^ K562 target cells (data from 2 independent donors). **h**, Quantification of CD69 surface expression (MFI) in Hybrid-R T cells 48 hours after stimulation with CD19^+^ K562 target cells (data from 2 independent donors). **i**, Quantification of IL-2 production in cell culture supernatant after coculture for 48 hours with CD19^+^ K562 target cells (data from 2 independent donors). Mean SEM depicted. **j**, Transcriptional circuit regulation under SNIPR or optimized Hybrid-R in primary human T cells (representative of 3 experiments).

**Extended data fig. 2:**
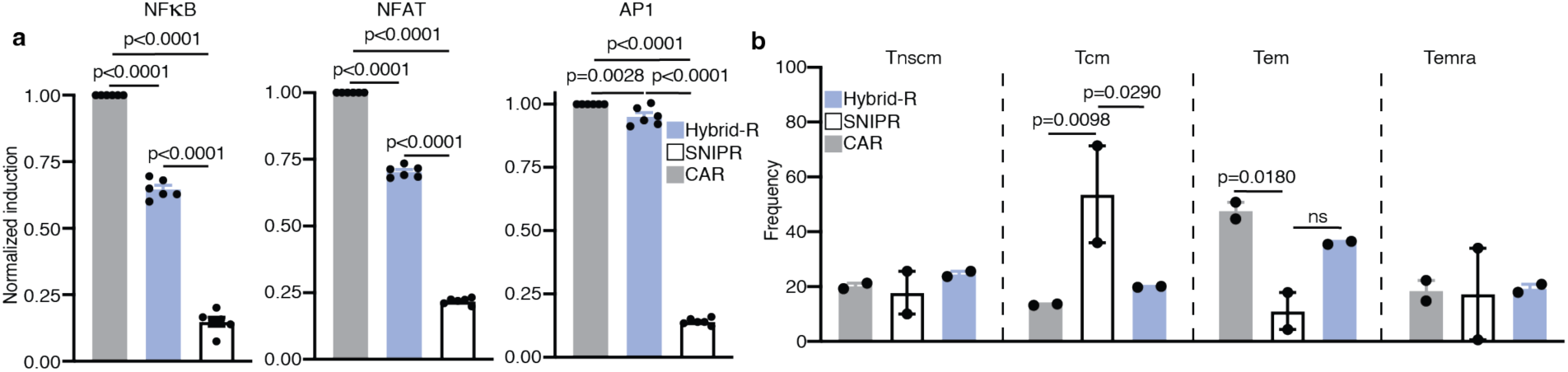
Direct comparison of optimized Hybrid-R, CAR, and SNIPR signaling. **a**, Normalized individual reporter activity (frequency) for CAR, SNIPR or Hybrid-R in Jurkat reporter T cells after 24-hour coculture with CD19^+^ K562 target cells (n=6 technical replicates). **b**, Quantification of T cell memory phenotype (by CD62L and CD45RA expression) in Hybrid-R T cells 48 hours after stimulation with CD19^+^ K562 target cells (data from 2 independent donors). Statistics calculated Statistics calculated with two-way ANOVA with Šídák’s multiple comparisons test.

**Extended data fig. 3:**
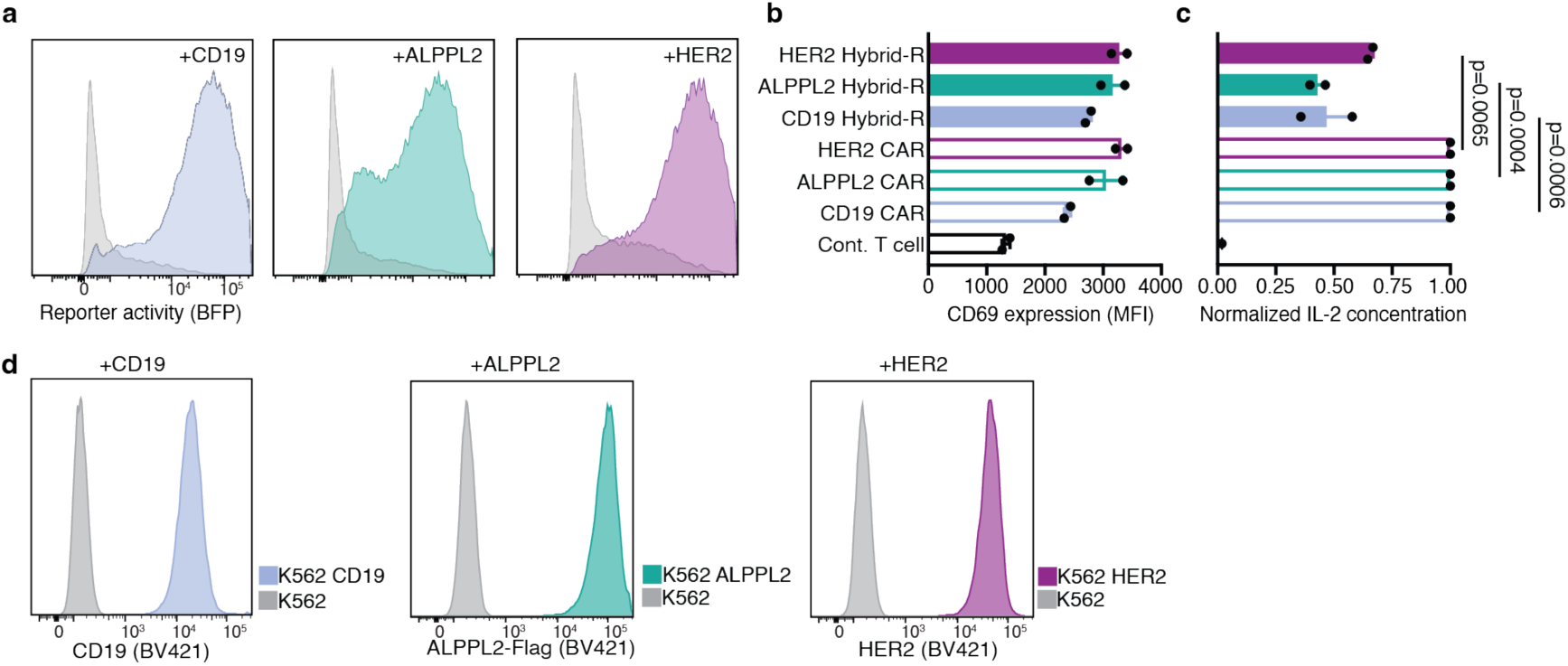
Hybrid-R senses and responds to multiple tumor associated antigens. **a**, BFP reporter gene expression in Hybrid-R primary human T cells 48 hours after stimulation with antigen^+^ K562 target cells (representative of 3 experiments). **b**, Quantification of CD69 surface expression (MFI) in Hybrid-R T cells 48 hours after stimulation with CD19^+^ K562 target cells. **c**, Normalized quantification of IL-2 production in cell culture supernatant is given after coculture for 48 hours with CD19^+^ K562 target cells. (data from 2 independent donors). Mean SEM depicted. **d**, Histograms depict target antigen expression on K562 cell lines.

**Extended data fig. 4:**
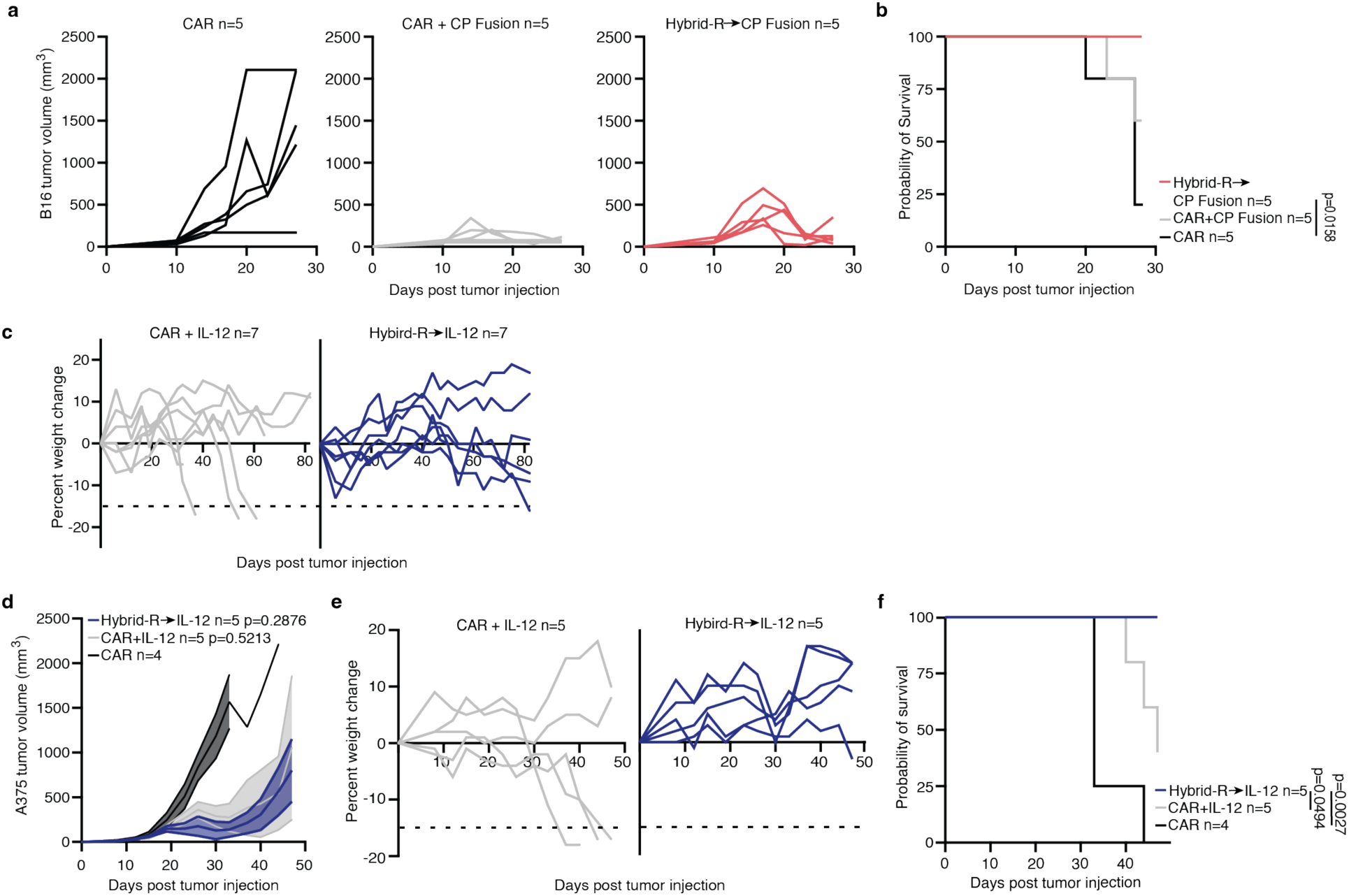
Hybrid-R regulation of T cell intrinsic and extrinsic payloads improves T cell therapy efficacy and safety across tumor models and in vivo backgrounds. **a**, Tumor volumes of hCD19^+^ B16 tumor-bearing animals treated with 4.0e6 engineered mouse T cells 11 days after tumor innoculation. Lines indicated individual mice. **b**, Survival of hCD19^+^ B16 tumor bearing syngeneic animals. P values calculated using log-rank Mantel-Cox (survival). **c**, CD19^+^ A549 tumor bearing NSG animal percent weight change. Lines indicate individual mice, 15% weight loss represents euthanasia criteria. **d**, Tumor volume from CD19^+^ A375 solid tumor-bearing NSG animals treated with 0.75e6 engineered T cells 10 days after tumor innoculation. Statistics analyzed using one-way ANOVA with Dunnett’s correction comparing the area under the curve versus the CAR control group over the course of the experiment. Mean SEM depicted. **e**, CD19^+^ A375 tumor NSG bearing animal percent weight change. Lines indicate individual mice, 15% weight loss represents euthanasia criteria. **f**, Survival of CD19^+^ A375 tumor bearing animals. P values calculated using log-rank Mantel-Cox (survival).

**Extended data fig. 5:**
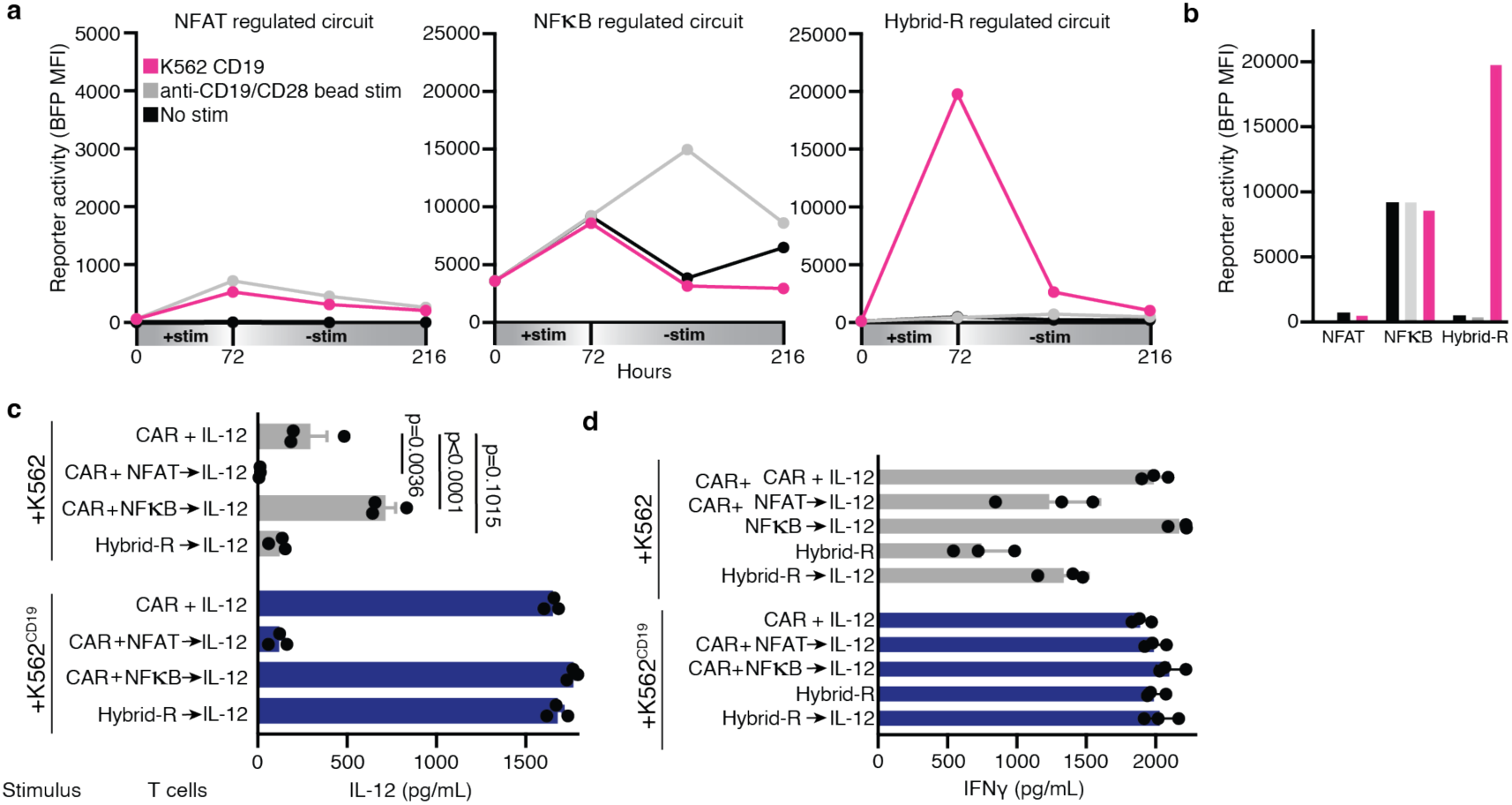
Investigation of Hybrid-R circuit kinetics and payload delivery capacity in vitro. **a**, Reporter circuit induction overtime from T cells engineered with anti-CD19 CAR and 4x NFAT or 5x NFkB response elements or Hybrid-R→BFP reporter circuit. Reporter MFI evaluated in presence or absence of stimuli (n=1 technical replicates). **b**, Reporter circuit MFI quantification at 72 hours after coculture (n=1 technical replicates). **c**, Quantification of human IL-12 production in cell culture supernatant is given after coculture for 72 hours with CD19^+^ K562 target cells (data from 3 independent donors). Statistics calculated with two-way ANOVA with Šídák’s multiple comparisons test. **d**, Quantification of human interferon-gamma production in cell culture supernatant is given after coculture for 72 hours with CD19^+^ K562 target cells for armored CARs and Hybrid-R→IL-12 T cells (n=3 technical replicates). Mean SEM depicted.

**Extended data fig. 6:**
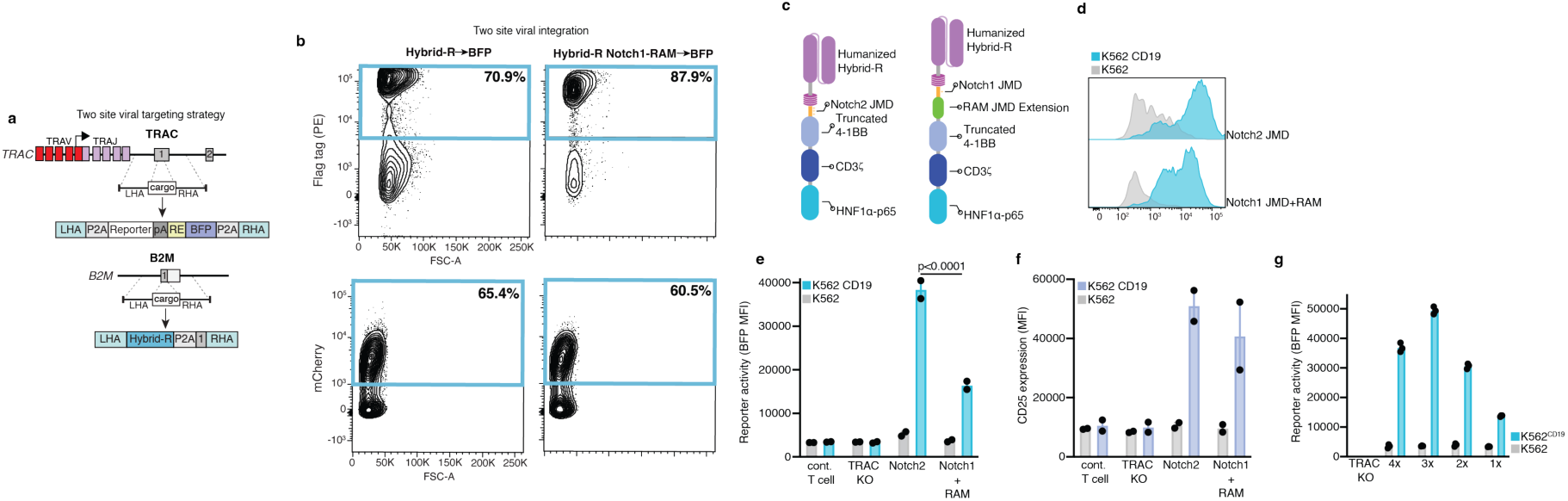
Humanization and site-specific integrated Hybrid-R circuits are active and tunable in primary human T cells. **a**, Schematic of two-site CRISPR/Cas9 + AAV viral Hybrid-R targeting strategy. **b**, Contour plots depict flag tag^+^ and mCherry^+^ primary human T cells at least 5 days after viral manufacutring. Gated on CD3^-^ /B2M^-^ (top) and CD45^+^ (bottom) cells. **c**, Schematic of humanized Hybrid-R designs. **d**, Histograms depict BFP reporter gene expression in Hybrid-R knock-in primary human T cells 72 hours after stimulation with target cells. **e**, Quantification (frequency of mCherry reporter^+^ cells) of BFP reporter expressing target cells after antigen encounter. Statistics calculated with two-way ANOVA with Šídák’s multiple comparisons test. **f**, Quantification (MFI) of CD25 surface expression on Hybrid-R T cells (data from 2 independent donors). **g**, Quantification (MFI) of BFP reporter expression after 72 hours with CD19^+^ K562 target cells.

**Extended data fig. 7:**
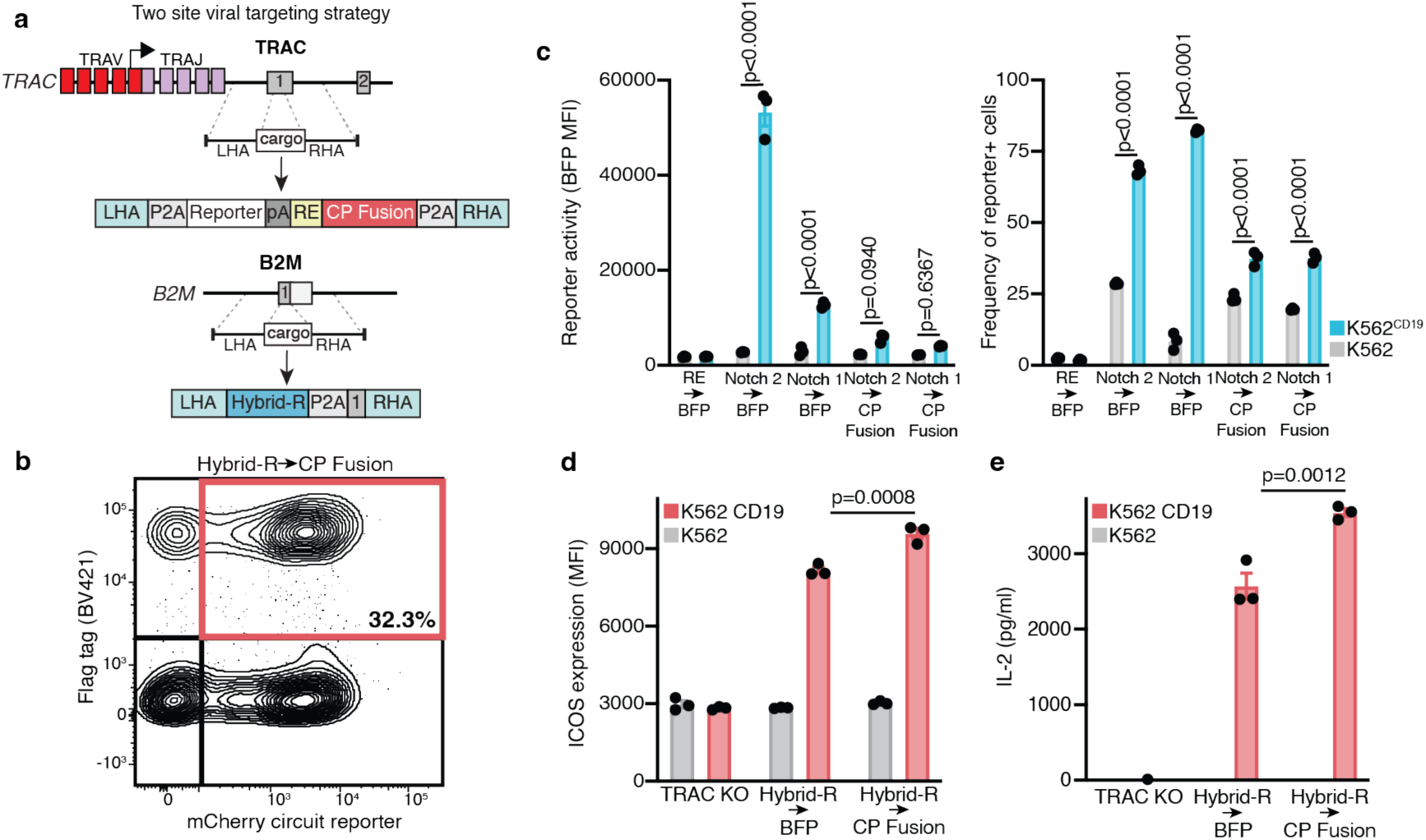
Two-site Hybrid-R→Payload circuits maintain enhanced T cell activity. **a**, Schematic of humanized Hybrid-R→CP Fusion T cell therapy targeting strategy. **b**, Contour plot depicts Flag tag^+^/mCherry^+^ T cells. Gated on CD3^-^/B2M^-^ cells. **c**, Left, Quantification (MFI) of BFP surface expression on Hybrid-R T cells. Right, Quantification (frequency of mCherry^+^ cells) of BFP reporter expression after 72 hours with CD19^+^ K562 target cells. Statistics calculated with two-way ANOVA with Šídák’s multiple comparisons test. **d**, Surface marker ICOS expression (MFI) on engineered T cells 72 hours after stimulation with CD19^+^ K562 target cells. **e**, Quantification of IL-2 in cell culture supernatant after 96 hour coculture with CD19^+^ K562 target cells via ELISA (n=3 technical replicates). Mean SEM is depicted.

**Extended data fig. 8:**
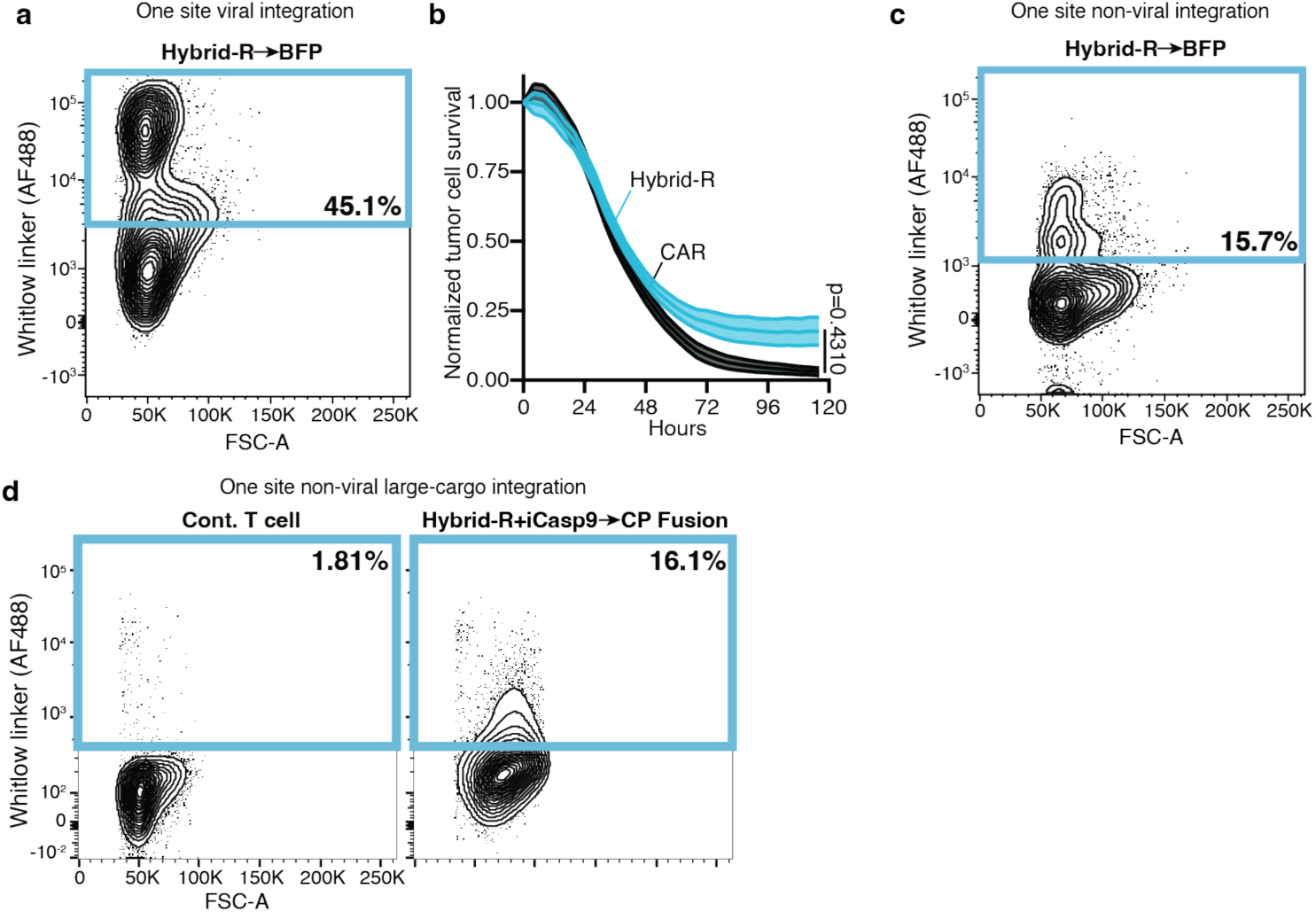
Single vector Hybrid-R activity in primary human T cells. **a**, Contour plots depict Whitlow linker^+^ primary human T cells at least 5 days after viral manufacutring. Gated on CD3^-^ cells. **b**, Relative target cell lysis by anti-B7-H3 Hybrid-Rs targeting B7-H3^+^ A549mKate-NLS cells measured by Incucyte live-cell imaging. Effector to target ratio is 1:4. Data are total area normalized to TRAC knockout T cells. Statistics were calculated using two tailed unparied T tests (data from 2 independent donors). Mean SEM depicted. **c**, Contour plots depict Whitlow linker^+^ primary human T cells at least 5 days after non-viral manufacutring. Gated on CD3^-^ cells. **d**, Contour plots depict Whitlow linker^+^ primary human T cells at least 5 days after non-viral manufacutring with clinical candidate Hybrid-R vector. Gated on CD3^-^ cells.

**Extended data fig. 9:**
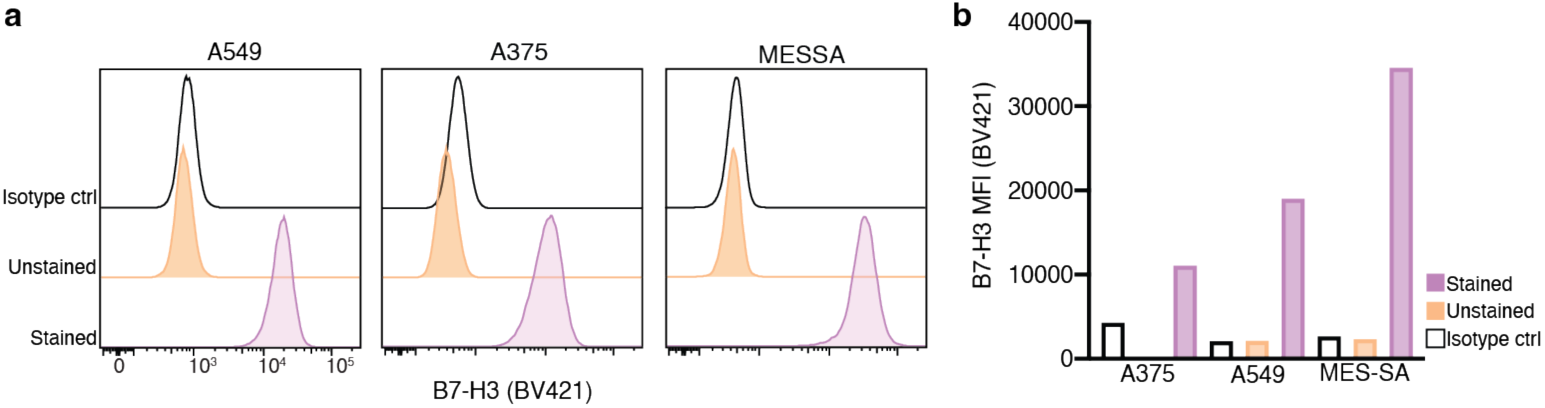
B7-H3 antigen density on solid tumor cell lines. **a**, Histograms depict B7-H3 surface expression across cell lines. **b**, Quantification (MFI) of B7-H3 surface expression across cell lines.

**Extended data fig. 10:**
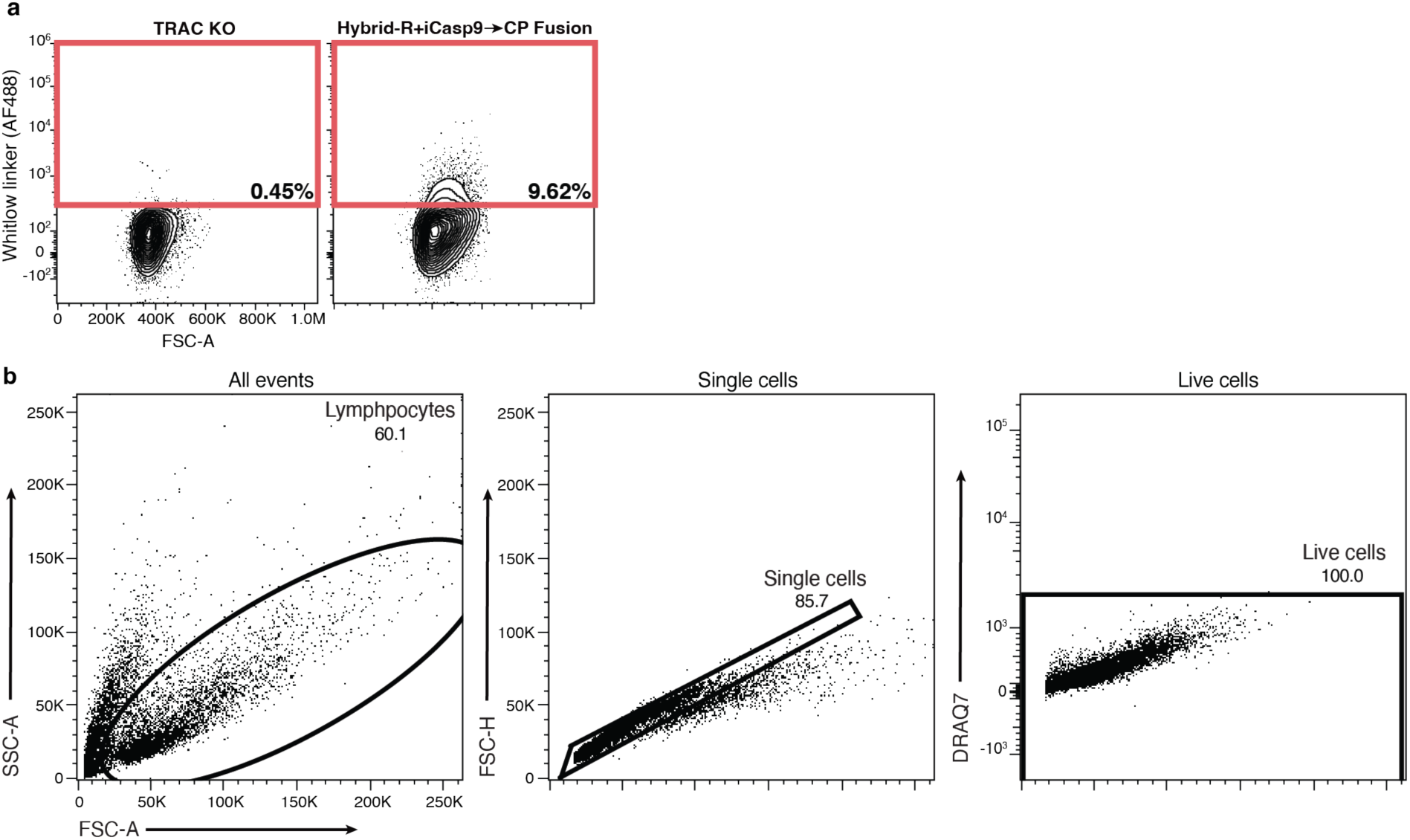
Clinical candidate Hybrid-R integration in primary human T cells. **a**, Contour plots depict clinical candidate Hybrid-R vector integration 10 days after electroporation. Gated on live cells. **b**, Example flow cytometry gating strategy for primary human T cell experiments.

## METHODS

### Receptor construction

Intracellular domains containing the appropriate costimulatory domain, CD3zeta domain, Gal4-VP64, HNF1α-P65 and GSlinkers were synthesized as gBlock gene fragments from Twist. Receptors were built by fusing the CD19 scFv or ALPPL2 scFv (M25^FYIA^) to the corresponding receptor scaffold and intracellular tail. All receptors contain an n-terminal CD8α signal peptide (MALPVTALLLPLALLLHAARP) for membrane targeting and a flag-tag (DYKDDDDK) or Whitlow linker (GSTSGSGKPGSGEGSTKG)^2^ for easy determination of surface expression with α-flag (Biolegend) or α-Whitlow (Cell Signaling). The receptors were cloned into a modified pHR’SIN:CSW vector containing a PGK or MSCV promoter for all lenti- and retroviral primary T cell experiments. Constitutive CP Fusion was cloned into the same pHR’SIN:CSW vector and was also fused to a T2A mCherry for easy determination of intracellular expression. The pHR’SIN:CSW vector was also modified to make the response element plasmids. Five copies of the Gal4 DNA binding domain target sequence (GGAGCACTGTCCTCCGAACG) or four copies of the HNF1α DNA binding domain target sequence (GTTAAT) were cloned 5′ to a minimal pybTATA promoter. Also included in lenti- or retroviral the response element plasmids is a PGK promoter that constitutively drives mCitrine expression to easily identify transduced T cells. For all inducible BFP, IL-12 and CP Fusion vectors, BFP, IL-12 or CP Fusion was cloned via a BamHI site in the multiple cloning site 3′ to the Gal4 or HNF1α response elements. All constructs were cloned via Infusion cloning (Clontech).

### Primary human T cell isolation and culture

Primary CD3+ T cells were isolated from anonymous donor blood after apheresis by negative selection (STEMCELL Technologies). Blood was obtained from Blood Centers of the Pacific, STEMCELL or Charles River as approved by the University Institutional Review Board. T cells were cryopreserved in RPMI-1640 (UCSF cell culture core) with 20% human AB serum (Valley Biomedical Inc.) and 10% DMSO. After thawing, T cells were cultured in human T cell medium consisting of either X-VIVO 15 (Lonza), 5% Human AB serum and 10 mM neutralized N-acetyl L-Cysteine (Sigma-Aldrich) supplemented with 30 units/mL IL-2 (PeproTech) or ImmunoCult-XF T Cell Expansion Medium (STEMCELL) supplemented with IL-7 and IL-15 (5 ng/mL each).

### AAV donor template cloning and AAV production

To generate pAAV-B2M donor vectors 500 bp homology arms flanking the Hybrid-R sequence were cloned into pAAV vector generously gifted by the Eyquem lab. To generate pAAV-TRAC donor vectors 300 pb homology arms flanking HNF1α response element including P2A-mCherry reporter, SV40 poly(A) sequence, HNF1α DNA binding domains and payload sequences were cloned into the pAAV vector generously gifted by Eyquem lab. To produce recombinant AAV6 (rAAV6) viruses, HEK293T cells were co-transfected with pAAV, pAAV6-Rep/Cap, and pAAV-Helper plasmids using the polyethylenimine (PEI) transfection protocol. 12 hours later the medium was replaced and three days later cell pellet and viral supernatant were collected. The cell pellet was suspended in cell lysis buffer containing 50 mM Tris HCL, 150 mM NaCl, 2 mM MgCl_2_, and lysed by four cycles of freeze-thaw in a dry ice/ethanol bath (10 minutes per cycle). The lysate was pelleted by spinning at 11,000 g for 10 minutes and the cleared supernatant was collected and treated with endonuclease Benzonase (Millipore) for 1 hour at 37°C. Viral supernatants were collected and precipitated using 40% PEG (Sigma) at 8% final concentration. Supernatant PEG mixture was shaken at 4°C for 1 hour and then kept at 4°C for 3 hours. Supernatant PEG mixtures were then spun down at 3800 rpm for 15 minutes at 4 °C, and the pellets were resuspended in cell lysis buffer and incubated with endonuclease Benzonase (Millipore) for 1 hour at 37°C. Lysates were combined and loaded into iodixanol gradient tubes and fractionated by ultra-centrifugation at 28,000 rpm for 16 hours at 6 °C. The 40% iodixanol layer was isolated using an 18 gauge needle and syringe. The supernatant was filtered and concentrated using a 50 K Amicon Ultra-15 centrifuge filter (Millipore). Concentrated rAAV6 viruses were aliquoted and stored at -80 °C for long-term storage. Titers were determined by real-time PCR using primers for the AAV inverted terminal repeat (ITR) sequence.

### sgRNA and RNP electroporation of primary human T cell and CRISPR knock-in

sgRNAs were designed using the CispRGold program and purchased from Integrated DNA Technologies (IDT)^1,2^. To produce sgRNA complexes equal volumes of 100 μM CRISPR-RNA (crRNA) and Trans-activating CRISPR RNA (tracrRNA) were mixed and heated at 95 °C for 5 minutes and cooled at room temperature. To produce RNP complexes, 100 pmol of cas9 protein was mixed with 200 pmol gRNA duplex. These mixtures were incubated for 10-20 minutes at room temperature for at least 10 minutes. 1.0x10^6^ primary human CD3+ T cells were suspended into 20μL of P3 electroporation buffer (Lonza) containing complexed RNPs. After electroporation with RNPs, T cells (EH100) were rescued with prewarmed ImmunoCult XF T Cell Expansion Medium (STEMCELL) supplemented with IL-7 and IL-15 5 ng/mL each. T cells were then transduced with rAAV6 donor particles at a multiplicity of infection (MOI) of 1-2x10^5^ genome copy per cell with 5 μM M3814 (MedChemExpress) DNA PK inhibitors and 5 μM ART558 (MedChemExpress) Polθ inhibitor. rAAV6 and inhibitors were removed 24 hours later. T cells were expanded and assessed for knock-in efficiency via flow cytometry.

### sgRNA and cas9 mRNA electroporation of primary human T cell and CRISPR knock-in

sgRNAs were designed using the CispRGold program and purchased from Integrated DNA Technologies (IDT)^1^. To produce sgRNA complexes equal volumes of 100 μM CRISPR-RNA (crRNA) and Trans-activating CRISPR RNA (tracrRNA) were mixed and heated at 95 °C for 5 minutes and cooled at room temperature. sgRNA complex was added to cas9 mRNA (400uM) and 500ng of HDRT nanoplasmid or HDRT PCR product. These mixtures were then added to 1.0x10^6^ primary human CD3+ T cells suspended in 20μL of P3 electroporation buffer (Lonza). After electroporation with mixture, T cells (EH115) were rescued with prewarmed ImmunoCult XF T Cell Expansion Medium (STEMCELL) supplemented with IL-7 and IL-15 5 ng/mL each. Rescued T cells were incubated for 30 minutes at 37 °C before plating with 5 μM M3814 (MedChemExpress) DNA PK inhibitors and 5 μM ART558 (MedChemExpress) Polθ inhibitor. T cells were expanded and assessed for knock-in efficiency via flow cytometry.

### GMP-compatible human Hybrid-R T cell therapy manufacturing

Primary CD3+ T cells were isolated from anonymous donor blood after apheresis by negative selection (STEMCELL Technologies). Isolated T cells were cultured in TheraPEAK X-VIVO15 media (Lonza) +5% Human AB Serum with IL-7 (20U/mL) and IL-15 (100U/mL) and activated with CTS CD3/CD28 Dynabeads (ThermoFisher) for 48 hours prior to electroporation. T cells were then collected and beads removed via magnetic separation. Combine duplexed sgRNA (320uM) with Cas9 (62.5uM) and incubate for 15 minutes at 37 °C. To the RNP mixture add the HDRT (Nanoplasmid sourced from Aldeveron) (15ug) in Genome Editing (GE) buffer (ThermoFisher) and incubate for 10 minutes at room temperature. Collect T cells and resuspend in GE buffer and mix with RNP, HDRT mixture. After electroporation T cells were rescued with prewarmed complete rescue media. Rescued T cells were incubated for 30 minutes at 37 °C before plating for expansion. T cells were expanded and assessed for knock-in efficiency via flow cytometry.

### Mouse T cell isolation culture and retroviral transduction

CD8^+^ T cells were isolated from the spleens of CD45.1^+^ or CD45.2^+^ mice using a mouse pan CD3^+^ T cell isolation kit (Biolegend). T cells were then cultured with 100 units/mL recombinant human IL-2 (PeproTech), T cells were stimulated overnight with anti-CD3 and anti-CD28 Dynabeads beads (ThermoFisher). Retroviral supernatants were added to T cells in plates coated with RetroNectin (Takara), and spin transduction was carried out for 1 hour at 2,000 rpm at 30 °C. Following transduction, T cells were resuspended and cultured in fresh medium containing 100 units/mL IL-2 until adoptive transfer. Transduction efficiency was determined by flow cytometry before adoptive transfer.

### Lentiviral transduction of human T cells

Pantropic VSV-G pseudotyped lentivirus was produced via transfection of Lenti-X 293T cells (Lenti-X 293T cells originate from female fetal tissue) (Clontech) with a pHR’SIN:CSW transgene expression vector and the viral packaging plasmids pCMVdR8.91 and pMD2.G using Mirus Bio TransIT-Lenti (Mirus). Primary T cells were thawed the same day, and after 24 hours in culture, were stimulated with Human T-Activator CD3/CD28 Dynabeads (ThermoFisher) at a 1:3 cell:bead ratio. At 48 hours, viral supernatant was harvested, and the primary T cells were exposed to the virus for 24 hours. At day 5 post T cell stimulation, the Dynabeads were removed, T cells were sorted and expanded until day 10-14 when they were rested and could be used *in vitro* or *in vivo* assays. T cells were sorted for assays with a Beckton Dickinson (BD) FACs ARIA II.

### Cancer cell lines

The cancer cell lines used were K562 (originating from female myelogenous leukemia cells) (ATCC), A549 (originating from male, lung epithelial carcinoma cells) (ATCC), A375 malignant melanoma (originally obtained from Dr. Alexander Marson’s laboratory at UCSF, originating from female melanoma cells), M28 human epithelioid cells (originally obtained from Dr. Brenda Gerwin’s laboratory at the National Cancer Institute), MES-SA (uterine sarcoma) (ATCC), and B16-F10 melanoma cells (ATCC). K562s, A375s and A549s were lentivirally transduced to stably express human CD19. B16-F10s were engineered to express human CD19. CD19 levels were determined by staining the cells with α-CD19 BV421 (Biolegend).

### *In Vitro* primary T cell assays

For all in vitro co-culture assays, purified primary human T cells transduced with Hybrid-R or Hybrid-R and circuit vectors were seeded with target cells at a 1:1 effector:target ratio for 48, 72 or 96 hours after which supernatants were isolated and frozen at -80 °C. Isolated mouse T cells transduced with Hybrid-R or Hybrid-R and circuit vectors were seeded with target cells at a 1:1 effector:target ratio for 96 hours after which supernatants were isolated and frozen at -80 °C. All IL-2 ELISA experiments (ThermoFisher) were conducted in medium without exogenous IL-2. Once culture supernatants were harvested, the remaining cells were stained for flow cytometric analysis for surface protein expression. All IL-12 ELISA experiments (RND Systems) were conducted in medium without exogenous IL-2. Once culture supernatants were harvested, the remaining cells were stained for flow cytometric analysis for surface protein expression. For proliferation assays T cells were stained with CellTrace far red proliferation dye following manufacturers protocol (ThermoFisher). All in vitro experiments conducted with CRISPR knock-in T cells were conducted in medium (ImmunoCult XF T Cell Expansion Medium (STEMCELL) without exogenous cytokines. After seeding, plates were spun at 200 *g* for 1 minute to promote interaction of CAR T cells and target cells.

### Jurkat reporter assays

Hybrid-R, CAR or SNIPR triple-reporter Jurkat cells (Jurkat T cells originate from a male T cell leukemia patient) were transduced in 12-well plate format with individual receptor lentiviruses. At 72 hours post transduction, cells were plated at a 1:1 effector:target ratio with CD19^+^ K562 or K562 cells. After seeding plates were spun at 200 *g* for 1 minute to promote interaction of CAR T cells and target cells.

### Flow Cytometry

For all in vitro primary T cell assays, cells were washed with PBS 2% FBS twice, stained with surface staining markers (diluted 1:200) in the dark at room temperature for 20 minutes, washed twice and resuspended in PBS 2% FBS with or without DRAQ7 (diluted 1:1,000). Some experiments utilize Zombie UV fixable viability Kit (Biolegend) following the manufacturers protocol. Cells were then analyzed on a Becton Dickinson FACSymphony X-50 flow cytometer or Becton Dickinson Fortessa.

### Incucyte Killing Assay

Target cell lines expressing nuclear mKate2 were seeded in a 96-well flat bottom plate. 24 hours later engineered T cells were added at an expected effector:target ratio (see Fig. details for exact ratios). Plates were imaged regularly using the IncuCyte S3 Live-Cell Analysis System (Essen Bioscience). Three-Five images per well were collected at x10 magnification. All IncuCyte experiments were performed in human T cell medium unless otherwise specified.

### *In Vivo* M28 Study

NOD.Cg-Prkdc^scid^ Il2rg^tm1Wjl^/SzJ (NSG) (UCSF LARC Breeding Core) female mice were dosed with 4.0 × 10^6^ ALPPL2^+^ M28 cells via flank subcutaneous injection. 7 days post tumor injection, Hybrid-R or CAR transduced T cells were dosed to tumor bearing animals via retro-orbital injection (see Fig. details for the number of T cells dosed per experiment). Mice with similar-sized tumors were randomized to receive treatments of engineered T cells. Digital caliper measurements were performed at regular timepoints to assess tumor volume. Tumor volume was calculated using the following formula: (length × width^2^)/2. Throughout experiment animal drinking water was supplemented with Clavomox (Zoetis) to prevent bacterial infections. All experimentation was performed in accordance with the IACUC guidelines present at UCSF.

### In Vivo A549 Study

NOD.Cg-Prkdc^scid^ Il2rg^tm1Wjl^/SzJ (NSG) (UCSF LARC Breeding Core) female mice were dosed with 1.0 × 10^6^CD19^+^ A549 (ATCC) via subcutaneous injection. 7 days post tumor injection, Hybrid-R or CAR transduced T cells were dosed to tumor bearing animals via retro-orbital injection (see Fig. details for the number of T cells dosed per experiment). Digital caliper measurements were performed at regular timepoints to assess tumor volume. Tumor volume was calculated using the following formula: (length × width^2^)/2. Throughout experiment animal drinking water was supplemented with Clavomox (Zoetis) to prevent bacterial infections. All experimentation was performed in accordance with the IACUC guidelines present at UCSF.

### *In Vivo* A375 Study

NOD.Cg-Prkdc^scid^ Il2rg^tm1Wjl^/SzJ (NSG) (UCSF LARC Breeding Core) female mice were dosed with 0.5 × 10^6^ CD19^+^ A375 cells via flank subcutaneous injection. 9-10 days post tumor injection, Hybrid-R or CAR transduced T cells were dosed to tumor bearing animals via retro-orbital injection (see Fig.s details for the number of T cells dosed per experiment). Mice with similar-sized tumors were randomized to receive treatments of engineered T cells. Digital caliper measurements were performed at regular timepoints to assess tumor volume. Tumor volume was calculated using the following formula: (length × width^2^)/2. Throughout experiment animal drinking water was supplemented with Clavomox (Zoetis) to prevent bacterial infections. All experimentation was performed in accordance with the IACUC guidelines present at UCSF.

### *In Vivo* Nalm6 Study

NOD.Cg-Prkdc^scid^ Il2rg^tm1Wjl^/SzJ (NSG) (UCSF LARC Breeding Core) female mice were dosed with 0.5 × 10^6^ Luciferase expressing Nalm6 cells via tail vein injection. 4 days post tumor injection, Hybrid-R or CAR transduced or knock-in T cells were dosed to tumor bearing animals via retro-orbital or tail vein injection (see Fig.s details for the number of T cells dosed per experiment). Bioluminescence imaging was performed using an IVIS Spectrum In Vivo Imaging system at regular timepoints to assess tumor burden. Animals were dosed with 200μL of 15mg/mL Luciferin via IP injection and allowed to ambulate for 12-20 minutes prior to capturing prone and supine images. Image capture time was adjusted based on bioluminescence intensity, and average radiance [p/s/cm²/sr] was used as a measurement of tumor burden. Throughout experiment animal drinking water was supplemented with Clavomox (Zoetis) to prevent bacterial infections. All experimentation was performed in accordance with the IACUC guidelines present at UCSF.

### *In Vivo* MES-SA Study

NOD.Cg-Prkdc^scid^ Il2rg^tm1Wjl^/SzJ (NSG) (UCSF LARC Breeding Core) mice were dosed with 4.0 × 10^6^ or 2.0 × 10^6^ B7-H3 MES-SA cells via flank subcutaneous injection. 5 or 8 days post tumor injection, Hybrid-R engineered T cells were dosed to tumor bearing animals via retro-orbital injection. Mice with similar-sized tumors were randomized to receive treatments of engineered T cells. Digital caliper measurements were performed at regular timepoints to assess tumor volume. Tumor volume was calculated using the following formula: (length × width^2^)/2. Throughout experiment animal drinking water was supplemented with Clavomox (Zoetis) to prevent bacterial infections. All experimentation was performed in accordance with the IACUC guidelines present at UCSF.

### *In Vivo* B16 Study

B6.SJL-Ptprca Pepcb/BoyJ female mice aged 6–12 weeks were injected subcutaneously with 1.0 × 10^5^ CD19-B16 melanoma cells. Mice with similar-sized tumors were randomized to receive treatments of engineered T cells. 9-11 days post tumor inoculation Hybrid-R or CAR transduced T cells were dosed to tumor bearing animals via retro-orbital injection. All animals received a dose of 2.0 × 10^6^ or 4.0 × 10^6^ receptor positive T cells (receptor positive frequency was assessed via flow cytometry on the day of injection) (see Fig. details for the number of T cells dosed per experiment). Digital caliper measurements were performed at regular timepoints to assess tumor volume. Tumor volume was calculated using the following formula: (length × width^2^)/2. All experimentation was performed in accordance with the IACUC guidelines present at UCSF.

### Statistics

Data points represent biological or technical replicates unless otherwise stated. Technical replicates were performed as distinct samples. No statistical methods were used to predetermine sample size.

